# rRNA intermediates coordinate the multilayered nucleolar phase transition in *C. elegans*

**DOI:** 10.1101/2023.01.23.525268

**Authors:** Demin Xu, Xiangyang Chen, Yan Kuang, Minjie Hong, Ting Xu, Ke Wang, Chuanhai Fu, Ke Ruan, Chengming Zhu, Xuezhu Feng, Shouhong Guang

**Affiliations:** Department of Obstetrics and Gynecology, The First Affiliated Hospital of USTC, The USTC RNA Institute, Ministry of Education Key Laboratory for Membraneless Organelles & Cellular Dynamics, School of Life Sciences, Division of Life Sciences and Medicine, Biomedical Sciences and Health Laboratory of Anhui Province, University of Science and Technology of China, Hefei, Anhui 230027, China; CAS Center for Excellence in Molecular Cell Science, Chinese Academy of Sciences, Hefei, Anhui 230027, P.R. China

**Keywords:** nucleolus, vacuole, ring, reshaping, rRNA, RNAP I, phase transition

## Abstract

The nucleolus is the most prominent membraneless organelle within the nucleus and plays essential roles in rRNA transcription and processing and ribosome assembly. How the structure of the nucleolus is maintained and regulated is poorly understood. Here, we identified two types of nucleoli in *C. elegans*. Type I nucleoli are spherical, and rRNA transcription and processing factors are evenly distributed throughout the nucleolus. In type II nucleoli, rRNA transcription and processing factors exclusively accumulate in the periphery rim, which is named the nucleolar ring. The hollow vacuole inside the nucleolar ring contains proteins that usually localize in the nucleoplasm but are capable of exchanging contents across the ring. The high-order structure of the nucleolus is dynamically regulated in *C. elegans*. Faithful rRNA processing is important to maintain the spherical structure of the nucleoli. The depletion of a class of rRNA processing factors, for example, class I ribosomal proteins of the large subunit (RPL), which are involved in 27SA_2_ rRNA processing, reshaped spherical nucleoli to a ring-shaped nucleolar structure. The inhibition of RNAP I transcription and depletion of two conserved nucleolar factors, nucleolin and fibrillarin, prohibits the formation of the nucleolar ring. We concluded that the integrity of nucleoli is highly dependent on rRNA processing and maturation, which may provide a mechanism to coordinate structure maintenance and gene expression.

## Introduction

The nucleolus is the most prominent membraneless organelle within the nucleus and forms around tandem arrays of ribosomal gene repeats, termed nucleolar organizer regions (1). The nucleolar proteome identified more than 1000 different proteins, most of which are involved in ribosome biogenesis, including rRNA transcription and processing and ribosome assembly (2, 3). An increasing number of studies suggest that the nucleolus is a multilayered biomolecular condensate that assembles via liquid– liquid phase separation (LLPS) (4–7). In vitro reconstitution studies suggested that the key nucleolar proteins, fibrillarin and nucleophosmin, can undergo LLPS (4). In *X. laevis* germinal vesicles, nucleoli exhibit liquid-like behavior and spontaneously coalesce and round up upon contact (5). In addition, it was suggested that the organization of nucleolar subcompartments arises through multiphase liquid immiscibility, which is driven by the differential surface tension of substructures (4, 8).

The nucleoli of mammalian cells display three internal phase-separated subcompartments, the fibrillar center (FC), the dense fibrillar component (DFC), and the granular component (GC) (7). Typically, the FC is surrounded by a shell of DFC; both are enclosed in the GC. Recent work has suggested that *C. elegans* nucleoli also contain two phase-separated subcompartments, GC and FC (9). In addition, there is a highly conserved central region, called the nucleolar cavity or vacuole, present in the nucleoli of various plants and animals, which was first observed in the 19^th^ century (Fig. 1A) (9–13). Similar vacuoles have been observed in mammalian cells, such as in MCF-7, COS-7 and Hep-2 cells (14).

**Figure 1.**
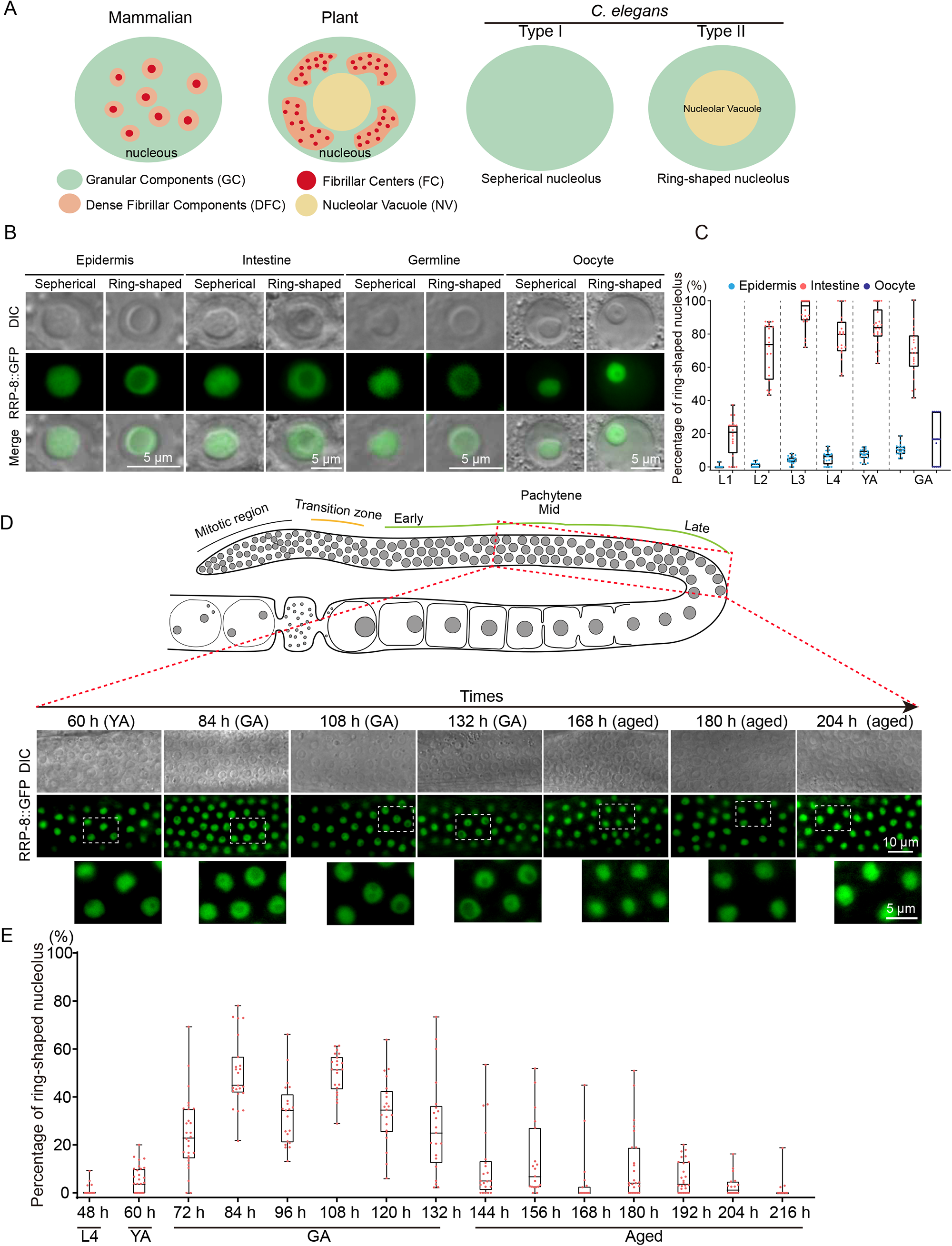
Identification of two-shaped nucleoli in *C. elegans*. (A) Schematic diagram of nucleolar structure in mammals, plants and *C. elegans*. (B) Differential interference contrast (DIC) and fluorescence microscopy images of *C. elegans* nucleoli in the indicated tissues. (C) Quantification of ring-shaped nucleoli at the indicated developmental stages. n>20 animals. (D) DIC and fluorescence microscopy images of pachytene cells. (E) Quantification of ring-shaped nucleoli in the germline pachytene stage. n>20 animals.

Although nucleolar vacuoles have been discovered for more than a hundred years, their regulatory mechanism, composition and function remain unclear. In *soybean,* different types of cellular stresses and environmental stimuli, for example, cold-warm treatment, led to the formation of nucleolar vacuoles (15, 16). Similarly, nucleolar vacuoles were formed in heat and chilling stress-treated *Arabidopsis thaliana* (17) and 5-fluoro-uracil-treated *Jerusalem artichoke tubers* (13). Drugs inhibiting rRNA synthesis, including actinomycin D, 5-fluoro-uracil and 2-thio-uracil, could impede vacuole formation in *tobacco callus* and *Zea mays* cells (18, 19). The vacuoles may contain ribonucleoprotein (RNP) complexes because both small nuclear RNAs (snRNAs) and small nucleolar RNAs (snoRNAs) have been detected (20, 21). It was postulated that nucleolar vacuoles may engage in mRNA surveillance and export, transport of nucleolar substances and temporary storage of certain materials (12, 13, 15, 22). In addition, larger vacuoles may be linked to higher nucleolar activity (15).

Here, we investigated the nucleolar structure of *C. elegans* and identified two types of nucleoli: spherical and ring-shaped nucleoli. We conducted candidate-based RNAi screening and identified a distinct class of RPLs, the knockdown of which reshaped spherical nucleoli to ring-shaped nucleoli and therefore were named class I RPLs. Interestingly, the ring-shaped nucleoli exhibited reentrant phase transition behavior by monotonically increasing 27SA_2_ pre-rRNAs. Through circularized reverse-transcription PCR (cRT-PCR), we detected abnormal accumulation of 27SA_2_ rRNAs upon the depletion of class I RPLs. The ring-shaped nucleoli had a subcompartment termed the nucleolar vacuole, which contains proteins usually localized in the nucleoplasm. NUCL-1 and FIB-1, two highly conserved nucleolar proteins with internal disordered sequences, were required for the formation of the nucleolar ring. The work suggested that the structure of the nucleolus is highly coordinated with rRNA processing and maturation.

## Results

### Identification of two-shaped nucleoli in *C. elegans*

During the culture of *C. elegans*, we frequently observed hollow nucleoli with rim and vacuole structures by differential interference contrast (DIC) microscopy (Fig. S1A). To further explore the nucleolar structure, we generated an RRP-8::GFP transgene to label the nucleoli (23). The nucleoli were visualized by DIC and fluorescence microscopy. We identified two types of nucleoli in *C. elegans*. In type I nucleoli, RRP-8::GFP likely evenly occupied the entire nucleoli, and the nucleoli were spherical under a DIC microscope (Fig. 1B). The type II nucleoli were convex in the periphery and concave in the center under DIC (Fig. 1B). RRP-8::GFP was strongly enriched in the periphery of the nucleoli and colocalized with the convex region but did not accumulate in the center. Therefore, we named the convex part the nucleolar ring and the concave part the nucleolar vacuole (Fig. 1A). Both types of nucleoli existed in different tissues (Figs. 1B-C). In epidermal cells and gonadal cells, the nucleoli were mainly spheres, and in intestinal cells, the nucleoli were mainly ring-shaped. The nucleolar structure in the germline is reshaped dynamically during development. The percentage of ring-shaped nucleoli in the germline was increased during germline maturity in young gravid adults and reduced in aged animals, suggesting reshaping processes between spherical and ring-shaped nucleoli (Figs. 1D-E).

### Class I *rpl* genes are required to maintain nucleolar structural integrity

To explore the mechanism of nucleolar structure formation, we conducted candidate-based RNAi screening to search for factors that are required to maintain the two-shaped nucleoli. We selected 110 genes that were involved in rRNA processing and ribosome assembly and knocked down these genes by feeding RNAi (Fig. 2A, Table S1). Then, we visualized the nucleolar morphology and localization of RRP-8::GFP. Of 48 *rpl* genes, knocking down a group of 19 *rpl* genes significantly increased the proportion of ring-shaped nucleoli (Figs. 2B-D, S1B-E). We divided the 48 *rpl* genes into two classes according to whether knocking down the gene could promote the formation of ring-shaped nucleoli (Fig. 2E). Knocking down class I *rpl* genes significantly increased the size of nucleoli and nucleolar vacuoles (Figs. 2F-H and S1F) and slightly increased the size of nucleolar rings (Figs. 2I-K).

**Figure 2.**
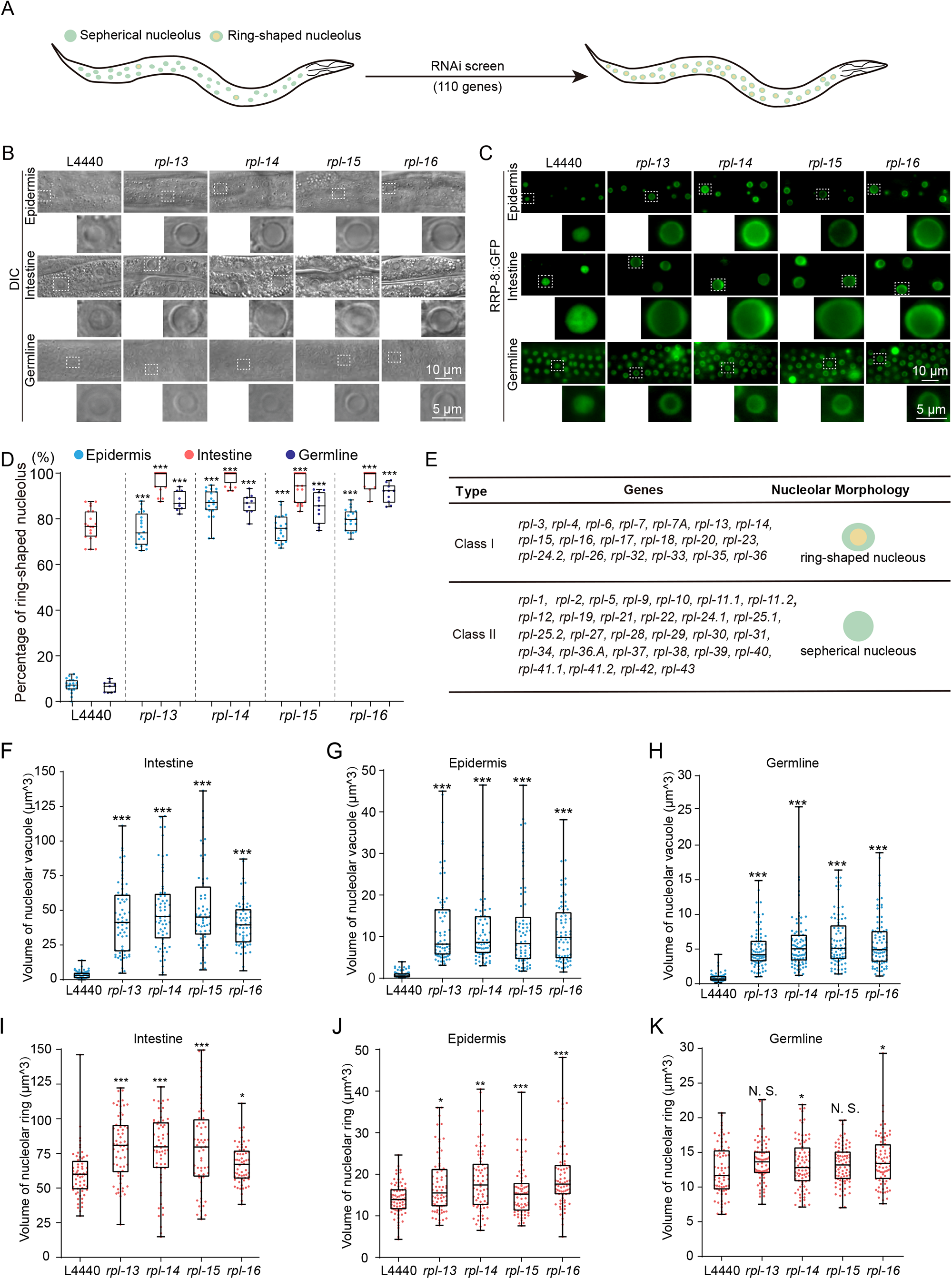
Class I RPLs are required for the maintenance of nucleolar structural integrity. (A) Schematic diagram of the candidate RNAi-based genetic screening for the formation of ring-shaped nucleoli in the *C. elegans* epidermis. (B-C) Images of *C. elegans* nucleoli after knockdown of the indicated genes by RNAi in distinct tissues. (D) Quantification of the ring-shaped nucleoli after RNAi knockdown of the indicated genes. n>11 animals. (E) Summary of the formation of ring-shaped nucleoli after knocking down Class I and II *rpl* genes. (F-K) The size of the nucleolar vacuole (F-H) and nucleolar ring (I-K) after RNAi knockdown of the indicated genes. n>60 nucleoli from 10 animals.

The nucleolus is likely formed by LLPS, which is modulated by environmental temperature. Many disordered proteins and/or nucleic acids undergo LLPS either upon cooling or heating (24–27). Heat and chilling stress have been shown to induce the formation of nucleolar vacuoles in *Arabidopsis thaliana* (17). Previously, we showed that cold stress induced the production of risiRNA and the accumulation of two nucleoplasmic proteins, NRDE-2 and NRDE-3, in the nucleoli in *C. elegans* (23, 28, 29). We then investigated the effect of cold stress on nucleolare reshaping. In wild-type animals, the ring-shaped nucleoli were suppressed by 4°C cold stress and reverted when shifting back to 20°C (Figs. S2A-B). Similarly, the formation of *rpl-14 knockdown*-induced ring-shaped nucleoli was strongly prohibited by cold stress, and this inhibition was released after transferring the worms to 20°C for 24 hours (Figs. S2C-D).

### rRNA transcription and processing machinery accumulate in the nucleolar ring

The factors involved in rRNA transcription and processing and ribosome assembly are largely evenly dispersed in the spherical nucleolus (23, 30). To test whether nucleolar reshaping could transform the localization of other rRNA transcription and processing factors in addition to RRP-8, we generated transgenes of a number of nucleolus-localized proteins and examined their localization upon knocking down *rpl-14* by RNAi. FIB-1 is a conserved fibrillarin involved in nucleologenesis and is usually used as a nucleolar marker (31, 32). RBD-1 is an ortholog of human RBM19, which is required for 90S pre-ribosome maturation (33). Both FIB-1 and RBD-1 occupied the entire spherical nucleolus in wild-type animals and accumulated in the nucleolar ring upon RNAi *rpl-14* (Figs. 3A-B). RPOA-1, RPOA-2 and RPAC-19 are subunits of the RNA polymerase I complex and are dispersed in the spherical nucleolus and accumulated in the nucleolar ring upon *rpl-14* RNAi (Figs. 3C-D, S3A). DAO-5, an rRNA transcription factor, also translocated to the nucleolar ring upon *rpl-14* RNAi (Fig. S3B). Consistently, DAO-5 has been previously reported to localize in the nucleolar ring in intestine cells in wild-type animals (31). rRNA processing proteins, such as RPL-7 and T06E6.1, were enriched in the nucleolar ring upon *rpl-14* RNAi (Figs. S3C-D). These results are summarized in Fig. 3E and imply that the nucleolar ring may still be capable of conducting rRNA transcription and processing.

**Figure 3.**
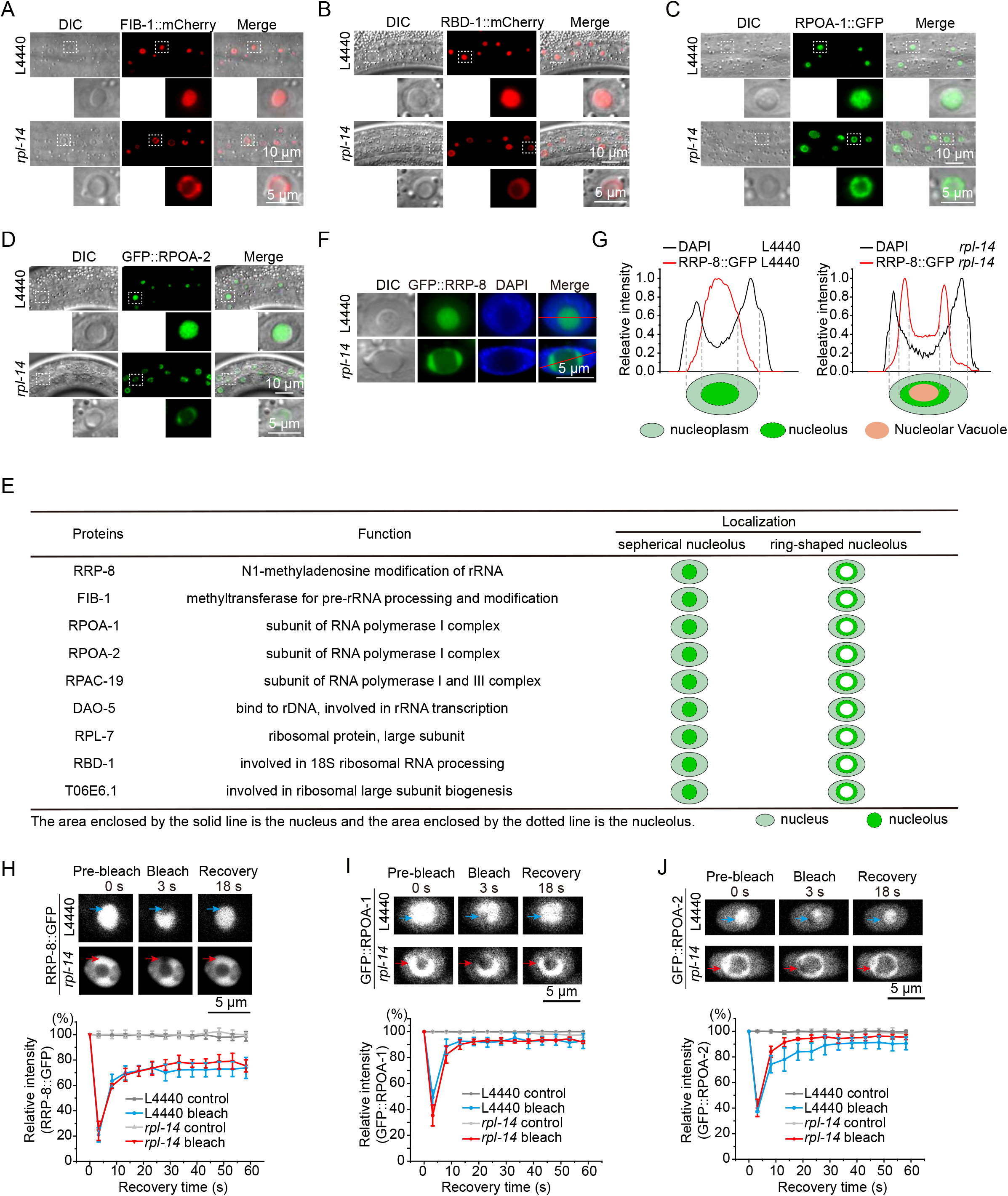
rRNA transcription and processing factors accumulate at the nucleolar ring. (A-D) DIC and fluorescence microscopy images of the indicated transgenes after knocking down *rpl-14*. (E) Summary of the localization of the indicated proteins after *rpl-14* RNAi. (F) DAPI staining of epidermal cells after *rpl-14* RNAi. (G) Fluorescent density scan of GFP:: RRP-8 and DAPI staining by ImageJ. (H-J) (top) FRAP assay of transgenes in the indicated regions before and after *rpl-14* RNAi. (bottom) Quantification of FRAP data. mean ± SD, n = 3.

We investigated the localization of nucleic acids after spherical to ring-shaped nucleolar reshaping. Both DAPI staining and GFP::HIS-71 (HIS-3.3) transgene indicated that DNA was enriched in the nucleoplasm in both spherical and ring-shaped nucleoli but likely depleted from the nucleolar vacuole (Figs. 3F-G, S3E-F). An RNA-specific dye, SYTO RNASelect, is usually used to stain abundant rRNAs. In control animals, RNA was evenly distributed in the whole spherical nucleoli (Fig. S3G-H). However, knocking down *rpl-14* enriched RNAs in the nucleolar ring but depleted them from nucleolar vacuoles (17).

The phase separation ability of nucleoli is essential for their function (7). We performed a FRAP assay to investigate whether nucleolar reshaping alters the mobility of rRNA transcription and processing machinery by comparing the mobility of RRP-8, RPOA-1 and RPOA-2 in spherical and ring-shaped nucleoli. All three proteins did not alter their mobility after bleaching in the sperical nucleoli and the nucleolar ring (Figs. 3H-J). These data suggested that nucleolar reshaping may not change the mobility of the components for rRNA transcription and processing and implied that the nucleolar ring is still able to conduct the reactions for rRNA transcription, processing and ribosome assembly.

### Nucleoplasmic proteins accumulate in nucleolar vacuoles

To investigate the composition of nucleolar vacuoles, we generated a number of fluorescence-labeled transgenes that usually accumulate in the nucleoplasm and are involved in pre-mRNA transcription and processing. We visualized their subcellular localization with and without *rpl-14* RNAi.

AMA-1 is the core subunit of RNA polymerase II. TAF-12 is a subunit of the transcription factor TFIID complex. GTF-2H2C is a subunit of the transcription factor TFIIH holo complex. AMA-1, TAF-12 and GTF-2H2C all accumulated in the nucleolar vacuole in wild-type animals (Fig. S4A) and upon *rpl-14* RNAi (Figs. 4A-C). MTR-4 and EMB-4 are two pre-mRNA processing factors and are involved in nuclear RNAi (34, 35). NRDE-2 and NRDE-3 are two proteins required for nuclear and nucleolar RNAi (28, 34, 36, 37). We observed vacuolar localization of these factors in both wild-type N2-background animals (Figs. S4B-D) and *rpl-14* RNAi animals (Figs. S4E-I). EXOS-2 is a subunit of the RNA exosome and accumulates in the nucleoplasm, nucleolar ring and vacuole (Fig. S4D). We summarized these data in Fig. 4D.

**Figure 4.**
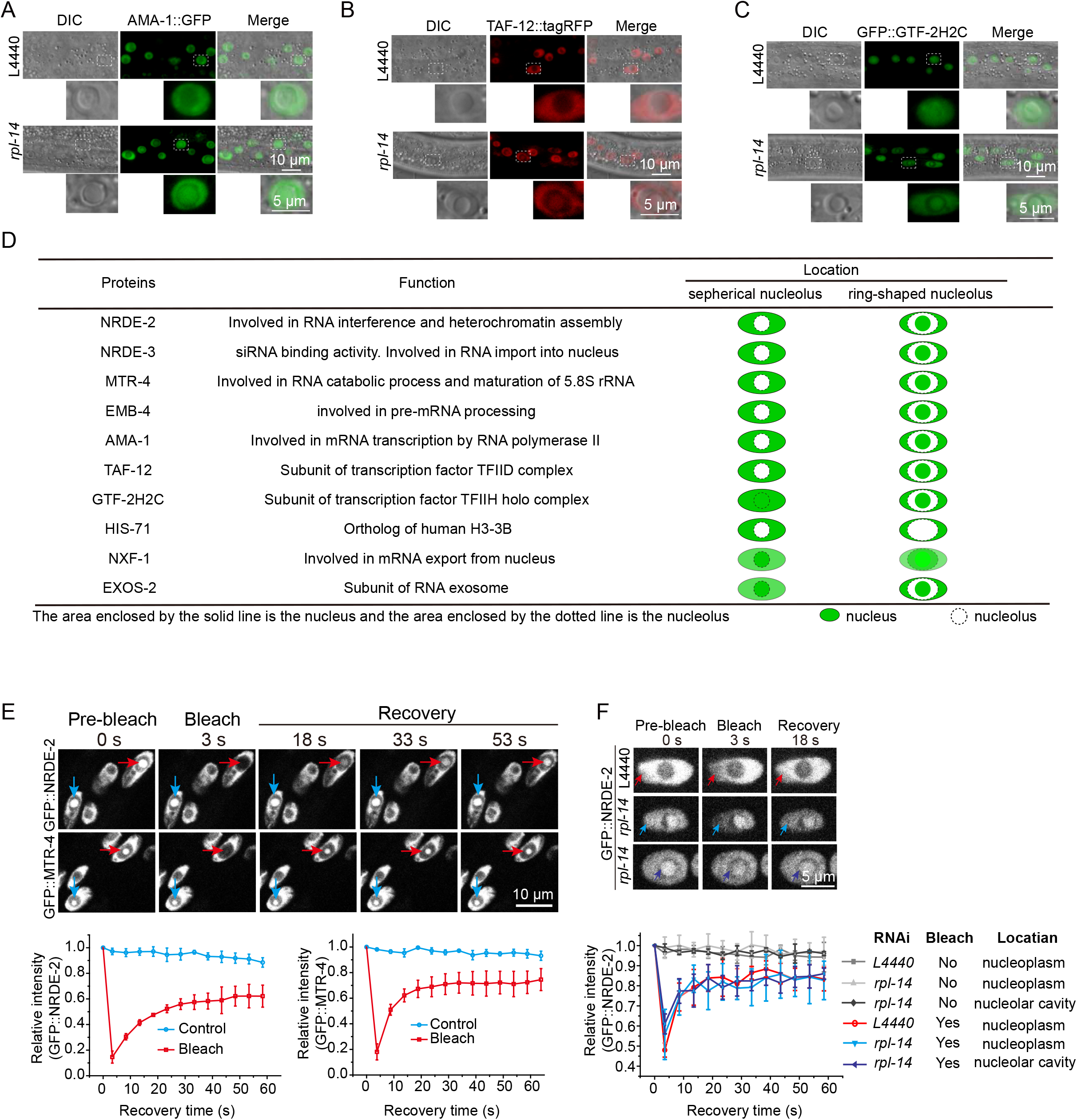
The nucleolar vacuole contains nucleoplasmic proteins. (A-C) DIC and fluorescence microscopy images of the indicated transgenes after knocking down *rpl-14*. (D) Summary of the localization of the indicated proteins after *rpl-14* RNAi. (E, F) (top) FRAP assay of GFP::NRDE-2 and GFP::MTR-4 in the indicated regions. (bottom) Quantification of FRAP data. mean ± SD, n = 3.

The nucleolus is physically separated from the nucleoplasm but rapidly exchanges its content with the surrounding nucleoplasm (7). To test whether the nucleolar ring could block the composition exchange between the nucleolar vacuole and nucleoplasm, we performed FRAP assays of GFP::NRDE-2 and GFP::MTR-4 in the nucleoplasm and nucleolar vacuole. Thirty seconds after bleaching, the GFP::NRDE-2 and GFP::MTR-4 fluorescence intensity in the nucleolar vacuole recovered to approximately 60% of the control fluorescent intensity (Fig. 4E). GFP::NRDE-2 exhibited similar mobility by FRAP in the nucleoplasm before and after *rpl-14* RNAi and in the nucleolar vacuole (Fig. 4F). However, GFP::NRDE-2 fluorescence did not recover when the whole nucleus was bleached (Fig. S4J). These data suggested frequent protein exchange between the nucleoplasm and the nucleolar vacuole.

NXF-1 is an mRNA export factor (38). NXF-1 was highly enriched in the nucleolar vacuole in wild-type N2 animals and upon *rpl-14* RNAi in epidermal cells (Figs. S5A-C). Interestingly, NXF-1 revealed a development-related vacuolar enhancement in germline cells in wild-type animals (Figs. S5D-E). In *A. thaliana,* similar vacuolar localization was reported for mRNA splicing proteins, U1 snRNP-specific proteins and exon‒exon junction complex proteins (39). Taken together, these data implied that the nucleolar vacuole is capable of conducting mRNA biogenesis and regulation.

### Class I *rpl* genes are required for 27SA_2_ rRNA processing

RPLs are proteins of the 60S ribosome subunit that are involved in ribosome assembly as well as pre-rRNA processing (40). To investigate the mechanism of nucleolar reshaping by class I RPL proteins, we adopted a circularized reverse-transcription PCR (cRT-PCR) method to analyze rRNA intermediates upon knocking down *rpl* genes (Fig. 5A) (41, 42). To validate the method, we performed cRT-PCR followed by sequencing the ends of mature 5.8S, 18S and 26S rRNAs (Fig. S6A-C). The results were largely consistent with sequences annotated by Wormbase (29, 43).

**Figure 5.**
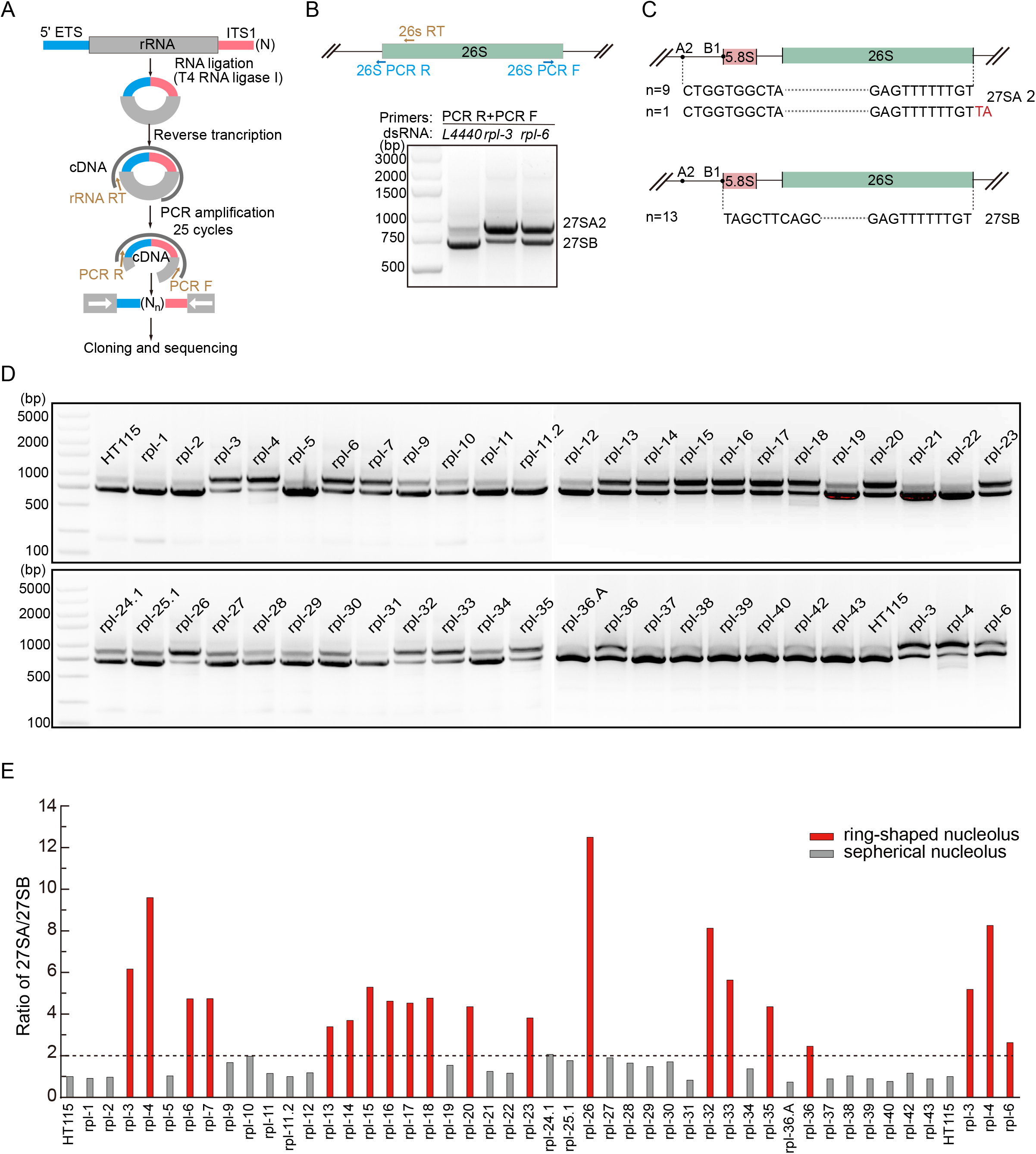
Class I *rpl* genes are required for 27SA_2_ rRNA processing. (A) Schematic diagram of circularized reverse-transcription PCR (cRT-PCR) method. (B) cRT-PCR assay of 26S pre-rRNA intermediates. (C) Sanger sequencing of the ends of 27SA_2_ and 27SB rRNAs. N represents the number of clones sequenced. (D-E) cRT-PCR assay (D) and quantification (E) of 27SA_2_ and 27SB pre-rRNA intermediates after knocking down the indicated *rpl* genes by RNAi.

Then, we assayed 26S rRNA intermediates by primer sets targeting the IVS II region upon knocking down *rpl* genes (Fig. 5B) and detected two major bands by the cRT-PCR method (Fig. 5B). We cloned and sequenced the 5’ and 3’ ends of these two bands and confirmed 27SA_2_ and 27SB pre-rRNAs (Fig. 5C). Strikingly, knocking down class I *rpl* but not class II *rpl* consistently led to the accumulation of 27SA_2_ pre-rRNA intermediates (Figs. 5D-E), implying that 27SA_2_ pre-rRNA or its processing may be involved in the formation of ring-shaped nucleoli.

RPS proteins are 40S ribosome subunits. We examined 18S rRNA intermediates after knocking down two *rps* genes, *rps-1* and *rps-5*. A number of additional 18S precursors were identified by cRT-PCR followed by sequencing (Figs. S7A-J). According to the identified 18S and 26S rRNA intermediates, a schematic diagram was drawn to reveal pre-rRNA processing steps (Fig. S8A).

To further confirm that 27SA_2_ rRNA is involved in the formation of ring-shaped nucleoli, we knocked down 26 predicted 60S rRNA processing factors by RNAi (Table S2). Feeding RNAi targeting 8 genes led to the formation of ring-shaped nucleoli (Figs. S8B-C). Most of the eight genes have been reported to be involved in 27SA and 27SB pre-rRNA processing (44, 45). Consistently, 27SA_2_ rRNAs accumulated upon knocking down these genes, as assayed by cRT-PCR (Figs. S7D-E).

Taken together, these data suggested that rRNA processing steps or intermediates were involved in the maintenance of nucleolar structure.

### FIB-1 and NUCL-1 are required for nucleolar reshaping

Intrinsically disordered regions (IDRs) are likely one of the driving forces of LLPS and have been identified in many phase-separated proteins (4, 46, 47). In addition, the GAR/RGG domain is an RNA binding segment that is frequently present in proteins capable of conducting LLPS (46–48). To further understand the mechanism of nucleolar reshaping, we searched the *C. elegans* genome for proteins that have both IDR and GAR/RGG domains and are predicted to localize in the nucleoli. We identified 8 proteins (Figs. 6A, S9A), among which knocking down *nucl-1* and *fib-1* strongly blocked the formation of *rpl-14*-induced ring-shaped nucleoli (Figs. 6B-C).

**Figure 6.**
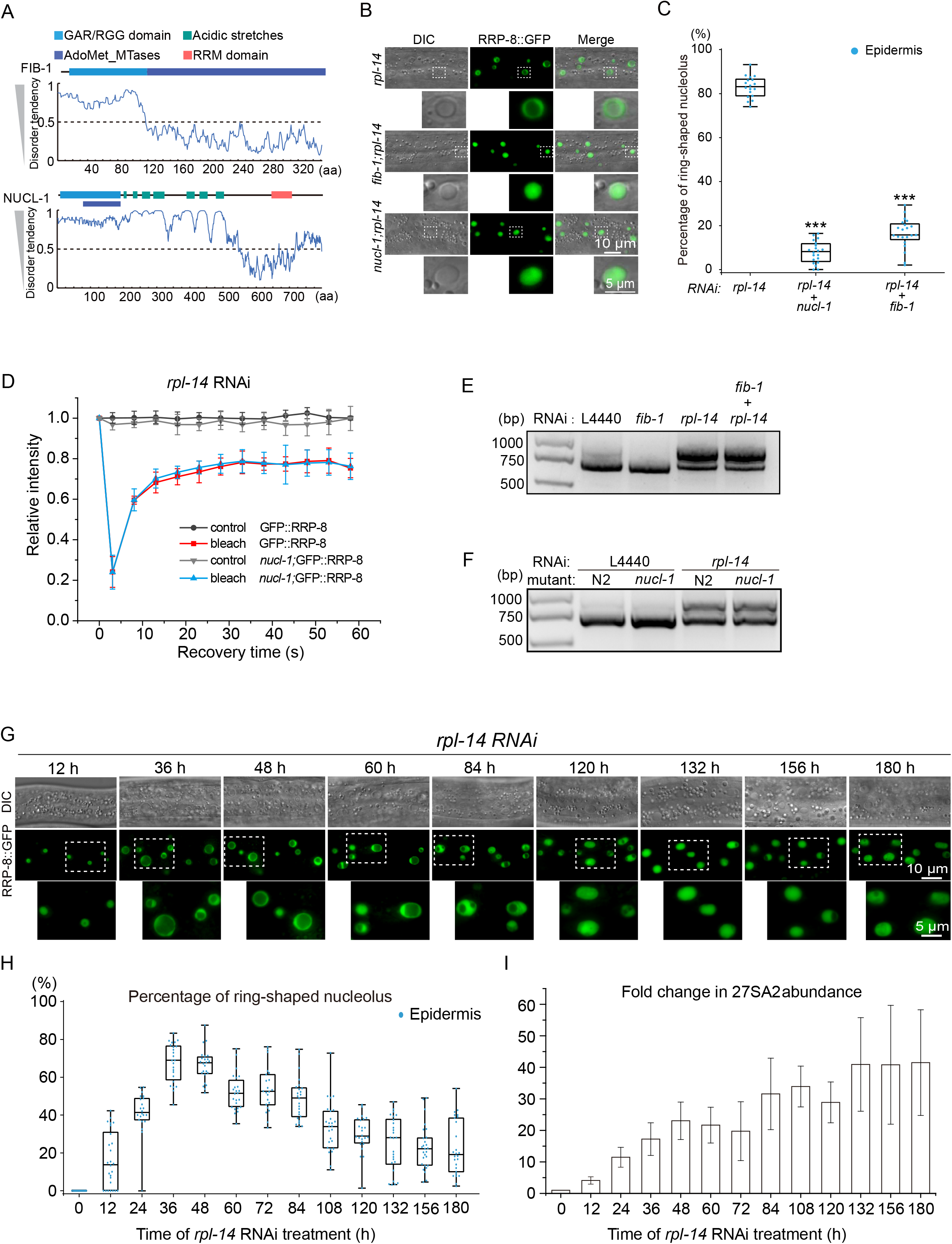
FIB-1 and NUCL-1 are required for the formation of ring-shaped nucleoli. (A) Schematic diagram of the domain structure and predicted intrinsically disordered regions of NUCL-1 and FIB-1. (B) DIC and fluorescence microscopy images of the indicated epidermal cells upon RNAi targeting of the indicated genes. (C) Quantification of ring-shaped nucleoli in epidermal cells. n=20 animals. (D) FRAP assay of GFP::RRP-8 in *nucl-1* mutants. mean ± SD, n = 3. (E) cRT-PCR assay of 27SA_2_ and 27SB pre-rRNA intermediates after knocking down the indicated genes by RNAi. (F) cRT-PCR assay of 27SA_2_ and 27SB pre-rRNA intermediates of the nucl-1 mutant after knocking down rpl-14 by RNAi. (G) DIC and fluorescence microscopy images of the indicated epidermal cells upon *rpl-14* RNAi over time. (H) Quantification of ring-shaped nucleoli in epidermal cells. n=20 animals. (I) Expression levels of 27SA_2_ pre-rRNAs quantified by real-time PCR.

FIB-1 is a highly conserved nucleolar protein with a GAR/RGG domain and methyltransferase domain that is involved in the methylation of pre-RNAs and nucleologenesis by LLPS (4, 22). NUCL-1 encodes an evolutionarily conserved protein exhibiting extensive homology to yeast and human nucleolin (9). The N-terminal domain of NUCL-1 is a long IDR containing a GAR/RGG domain, a methyltransferase domain, and an acidic stretch (Fig. 6A). The C-terminus of NUCL-1 harbors an RRM domain. Previous studies have shown that nucleolin associates with nascent pre-rRNA (49). Nucleolin in mammalian cells displays high mobility and is likely involved in phase separation (50). To confirm that NUCL-1 is required for the formation of ring-shaped nucleoli, we generated two additional deletion alleles of NUCL-1 by CRISPR/Cas9 technology (Fig. S9B). Both alleles inhibited the formation of *rpl-14 knockdown*-induced ring-shaped nucleoli (Figs. S9C-D). The FRAP assay revealed that the spherical nucleoli exhibited similar mobility in *nucl-1;rpl-14* animals to those in control animals (Fig. 6D). cRT-PCR showed that FIB-1 and NUCL-1 did not block the accumulation of 27SA_2_ (Figs. 6E-F, S9E-F), which suggested that FIB-1 and NUCL-1 functioned downstream of 27SA_2_ accumulation.

In vitro experiments showed that monotonically increasing RNA concentration could induce the formation of dynamic hollow condensates at high RNA-to-RNP ratios, but a disappearance at further higher RNA-to-RNP ratios, through multivalent heterotypic interactions that mediate a reentrant phase transition of RNPs containing arginine-rich IDR (51–53). To test whether nucleoli can undergo a reentrant phase transition in vivo during the nucleolar reshaping process, we conducated a time course of *rpl-14* RNAi and examined the nucleolar morphology. Surprisingly, knocking down *rpl-14* induced the formation of nucleolar rings during the early phase of RNAi, yet the ring-shaped nucleoli gradually disappeared after long-term *rpl-14* RNAi (Figs. 6G-H), suggesting an in vivo reentrant phase transition of the nucleoli (51). Consistently, 27SA2 rRNA monotonically increased during the time course of *rpl-14* RNAi (Figs. 6I, S9G).

Taken together, we speculate that NUCL-1 and FIB-1 participate in the regulation of nucleolar morphology through phase transition.

### Inhibiting rRNA transcription prohibits nucleolar reshaping

Nucleoli undergo dramatic changes when encountering cellular stresses and environmental stimuli. A previous study revealed that actinomycin D, which inhibits RNA polymerase I activity, could impede vacuole formation (18, 19).

To investigate the mechanism of the spherical-to-ring transition, we knocked down *rpl-14* by RNAi in the presence of actinomycin D. A 10 µg/ml actinomycin D treatment did not noticeably change the localization of RRP-8 or the size of nucleoli in wild-type N2 animals (Fig. 7A) but inhibited the formation of *rpl-14 knockdown*-induced nucleolar rings (Figs. 7B-C). At a higher concentration of actinomycin D (20 µg/ml), the *rpl-14*-induced nucleolar ring was completely inhibited, and RRP-8 likely accumulated as a nucleolar cap structure (54). Consistently, actinomycin D treatment inhibited the accumulation of *rpl-14*-induced 27SA_2_ rRNA intermediates by cRT-PCR assay (Figs. 7D).

**Figure 7.**
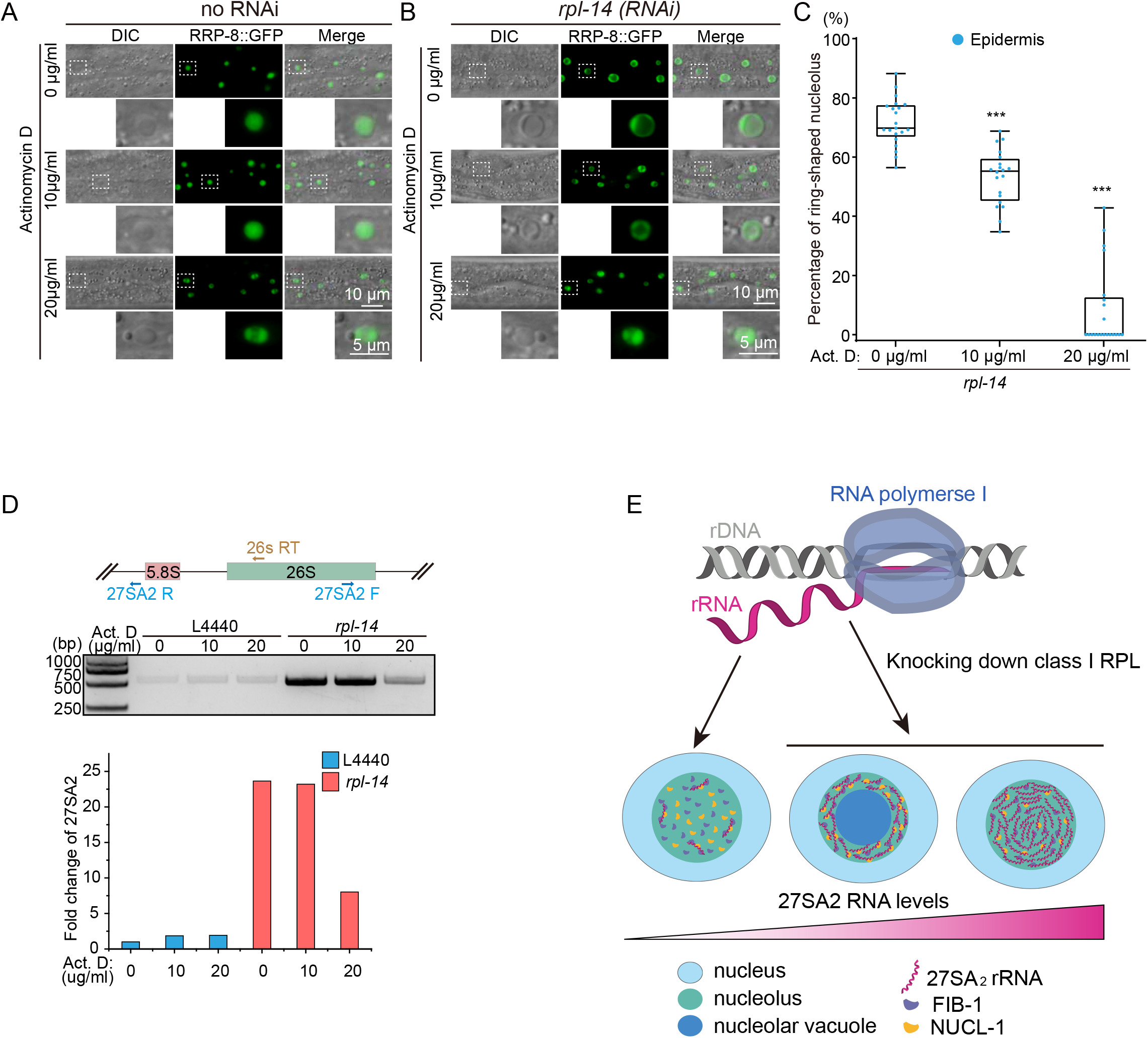
rRNA transcription and maturation are required for nucleolar reshaping. (A, B) DIC and fluorescence microscopy images of the indicated epidermal cells after actinomycin D (Act. D) treatment, without (A) or with (B) *rpl-14* RNAi. (C) Quantification of ring-shaped nucleoli after actinomycin D treatment in the presence of *rpl-14* RNAi. n = 20. (D) (top) cRT-PCR assay and (bottom) quantification of 27SA_2_ pre-rRNA intermediates after actinomycin D treatment in the presence of *rpl-14* RNAi. (J) A working model of rRNA intermediate-directed nucleolar reshaping.

## Discussion

The nucleolus is the most important membraneless organelle in the cell. Here, we observed two types of nucleoli, spherical and ring-shaped. The knockdown of class I RPL proteins, which are involved in 27SA_2_ rRNA processing, reshaped the spherical nucleoli to ring-shaped nucleoli (Fig. 7E). In ring-shaped nucleoli, rRNA transcription and processing factors accumulate in the nucleolar ring, and a large number of nucleoplasmic proteins accumulate in the nucleolar vacuole. The inhibition of RNAP I transcription by actinomycin D and depletion of two conserved nucleolar factors, nucleolin and fibrillarin, prohibit the formation of nucleolar rings.

### Regulation of nucleolar morphology by 27SA_2_ rRNA

The formation of multiphasic nucleolar structures may be due to the interaction of nucleolar proteins and rRNAs (4). Here, we found that the factors inhibiting the formation of ring-shaped nucleoli are all involved in 27SA_2_ rRNA processing, suggesting that the 27SA_2_ rRNA intermediates may play an important role in nucleolar morphology regulation. Similar to the ring-shaped nucleolus, [RGRGG]_5_ polypeptides and cellular RNAs can form hollow condensates in vitro (53). An in vitro experiment indicated that monotonically increasing the RNA concentration in mixtures of synthetic peptides containing multivalent arginine-rich linear motifs can induce dynamic droplet substructure formation and disapperance (51). Poly-L-lysine and single-stranded oligodeoxynucleotide droplets were shown to undergo repeated cycles of vacuole nucleation, growth and expulsion in applied electric fields (55). The RNA binding protein TDP-43 can also form hollow condensates in cells by LLPS if its RNA-binding capacity is disrupted (56). Here, we found that in *C. elegans*, the nucleoli exhibited similar reentrant phase transition behavior upon *rpl-14* RNAi, which can be inhibited by knocking down two conserved RNA binding proteins, NUCL-1 and FIB-1, both of which contain arginine-rich IDR. Further experiments are required to test the causal relationship between 27SA_2_ rRNAs and the formation of NUCL-1 or FIB-1 hollow condensates. Additionally, rRNA processing and maturation steps are highly conserved among eukaryotes. It will be very interesting to test whether 27SA_2_ rRNAs are involved in the maintenance of the hollow structure in plants or DFC/GC regions in mammalian cells.

It is unclear how and why 27SA_2_ rRNAs are involved in the formation and disappearance of ring-shaped nucleoli. We failed to observe nucleolar reshaping by injecting in vitro transcribed 27SA_2_ rRNAs into the *C. elegans* germline. It is unclear whether the in vitro transcribed rRNAs could diffuse into the nucleus or require certain nucleotide modifications. Previous work showed that N6-methyladenosine (m6A) modification of mRNAs can also induce m6A-binding proteins to undergo LLPS (57). Further investigations are required to investigate the role of 27SA2 rRNA in driving nucleolar reshaping in *C. elegans*.

### Phase transition of multilayered nucleolar condensates

According to the reentrant phase transition model, condensation is driven by electrostatic attraction between the negatively charged RNAs and the positively charged R/G-rich IDR polypeptide (51). However, excess negatively charged RNAs lead to the accumulation of a large number of counterions on the surface of the positively charged polypeptide, which triggers long-range electrostatic repulsion to prevent protein condensation (51, 53). An interplay between short-range cation−π attraction and long-range electrostatic repulsion may tune the reentrant phase transition (52). The charge balance of electrostatic interactions may be crucial for retaining transcriptional condensate in vivo (58). Based on these studies, we speculated that knocking down class I RPL may accumulate 27SA2 rRNAs and invert charges of nucleolar ribonucleoproteins, resulting in nucleolar reshaping.

Alternatively, nucleolar subcompartments may be formed by distinct protein droplets with different surface tensions (4, 8). For instance, HSP70 family proteins are enriched in the center of hollow condesation and retain the morphology of TDP43 spherical shells (56). The nucleolar vacuole of *C. elegans* was enriched with many nucleoplasmic proteins. Whether these nucleoplasmic proteins are involved in nucleolar reshaping is unclear. It will be very interesting to test whether the morphology of the ring-shaped nuceloli ring is maintained by certain specific proteins in the nucleolar vacuole or some disordered proteins on the nucleolar ring.

Previous reports have shown that the inhibition of nucleolar function can dramatically alter its structure (54, 59). Here, the inhibition of RNAP I by actinomycin D prohibited the formation of ring-shaped nucleoli, yet the mechanism is unclear. *C. elegans* exhibits different proportions of nucleolar ring structures in distinct tissues. Whether and how development and environmental stimuli reshape nucleolar structure require further investigation.

### The function of nucleolar vacuoles

The nucleolar vacuole is an evolutionarily conserved nucleolar subcompartment, but its function remains unclear. It has been speculated that the size of the vacuole represents nucleolar activities in plants (15, 16). We found that a certain proportion of nucleoli in various tissues in wild-type *C. elegans* were ring-shaped with nucleolar vacuoles inside. In intestinal cells, approximately 90% of the nucleoli have vacuole structures from larva to gravid adult. Of note, the nucleolar vacuole in the germline nucleoli revealed dynamic changes during aging. Interestingly, during germline development, an mRNA transporter, NXF-1, was specifically enriched in the nucleolar vacuole of the germline in gravid adult worms. Furthermore, the nucleolar vacuole was enriched with many nucleoplasmic proteins. Although the nucleolar ring barriers the vacuole with the nucleoplasm, there is rapid component exchange between these two compartments. These data suggested that nucleolar vacuoles may have important roles in germline development or mRNA metabolism. Consistently, previous studies have shown that nucleolar vacuoles may be involved in the transport of nucleolar substances from the nucleolus to the nucleoplasm and temporary storage of certain materials (12, 13, 22). In tobacco BY-2 cells, time-lapse photography revealed that the nucleolar vacuole slowly disappears over time (22). We also observed a dynamic change in nucleolar vacuoles over time in the oocyte nucleoli of *C. elegans*, suggesting that the vacuole is not a simple place for material storage but rather plays important regulatory roles.

## Materials and Methods

### Strains

Bristol strain N2 was used as the standard wild-type strain. All strains were grown at 20°C unless otherwise specified. The strains used in this study are listed in Supplementary Table S3.

### Candidate-based RNAi screening

RNAi experiments were performed at 20°C by placing synchronized embryos on feeding RNAi plates as previously described (60). HT115 bacteria expressing the empty vector L4440 (a gift from A. Fire) were used as controls. Bacterial clones expressing double-stranded RNAs (dsRNAs) were obtained from the Ahringer RNAi library (61) and sequenced to verify their identity. Some bacterial clones were constructed in this work, which are listed in Supplementary Table S4. All RNAi feeding experiments were performed for two generations except for larval arrest or sterile worms.

### Imaging

Images were collected using a Leica DM4 microscope.

### Construction of plasmids and transgenic strains

For ectopic transgenes, the promoter and CDS region and UTR were amplified from N2 genomic DNA. The mCherry coding sequence was amplified from PFCJ90. The vector fragment was PCR amplified from plasmid pSG274. These fragments were joined together by Gibson assembly to form the repair plasmid with the ClonExpress MultiS One Step Cloning Kit (Vazyme Biotech, Nanjing, China, Cat. No. C113-01/02). The transgene was integrated into *C. elegans* chromosome II via a modified counterselection (cs)-CRISPR method. The sequences of the primers are listed in Supplementary Table S5.

For in situ knock-in transgenes, the 3xFLAG::GFP coding region was PCR amplified from shg1248 genomic DNA. The GFP::3xFLAG coding region was PCR amplified from shg2123 genomic DNA. The mCherry coding region was PCR amplified from shg1660 genomic DNA. The tagRFP coding region was PCR amplified from YY1446 genomic DNA. Homologous left and right arms (1.5 kb) were PCR amplified from N2 genomic DNA. The backbone was PCR amplified from the plasmid pCFJ151. All these fragments were joined together by Gibson assembly to form the repair plasmid with the ClonExpress MultiS One Step Cloning Kit (Vazyme Biotech, Nanjing, China, Cat. No. C113-01/02). This plasmid was coinjected into N2 animals with three sgRNA expression vectors, 5 ng/µl pCFJ90 and the Cas9 II-expressing plasmid. The sequences of the primers are listed in Supplementary Table S6.

### CRISPR/Cas9-mediated gene editing

For the *nucl-1(ust313)* in-frame mutant, a 1.5 kb homologous left arm was PCR amplified with the primers 5′-GGGTAACGCCAGCACGTGTGGCCAAAGTTTAATCACCTCGCTCGC-3’ and 5’-TCGCTAAAACCAACTCGGCTTGAGTCGAAACCCATTTTGATTGTACC-3’. A 1 kb homologous right arm was PCR amplified with the primers 5′-AGCCGAGTTGGTTTTAGCGATAAGAGAAAACAGTATGATAG-3’ and 5’-CAGCGGATAACAATTTCACATCATCTTCATCCTCATCGTC-3’. The backbone was PCR amplified from the plasmid pCFJ151 with the primers 5′-CACACGTGCTGGCGTTACCC-3′ and 5′-TGTGAAATTGTTATCCGCTGG-3′. All these fragments were joined together by Gibson assembly to form the *nucl-1* plasmid with the ClonExpress MultiS One Step Cloning Kit (Vazyme Biotech, Nanjing, China, Cat. No. C113-01/02). This plasmid was coinjected into N2 animals with three sgRNA expression vectors, *nucl-1* sgRNA#1/#2/#3, 5 ng/µl pCFJ90 and Cas9 II expressing plasmid.

The sgRNAs used in this study for transgene construction are listed in Supplementary Table S7.

### Actinomycin D treatment

Actinomycin D (MedChemExpress no. HY-17559, CAS:50-76–0) was prepared at 20 mg/ml in DMSO as a stock solution. Each 3.5 µl actinomycin D stock solution was diluted with 300 µl Luria-Bertani liquid medium and layered onto NGM and RNAi plates. NGM and RNAi plates were prepared and placed at 37°C overnight before use. Synchronized L1 worms were placed onto the seeded plates and grown for 48 h before imaging and collection for cRT-PCR.

### Fluorescence recovery after photobleaching (FRAP)

FRAP experiments were performed using a Zeiss LSM980 laser scanning confocal microscope at room temperature. Worms were anesthetized with 2 mM levamisole. A region of interest was bleached with 100% laser power for 3-4 seconds, and the fluorescence intensities in these regions were collected every 5 s and normalized to the initial intensity before bleaching. For analysis, image intensity was measured by Mean and further analyzed by Origin software.

### DAPI staining

DNA was stained with DAPI Staining Solution (10 µg/ml) (Biosharp, BL105A) at room temperature. Worms were fixed with 1% formaldehyde before staining and soaked in DAPI solution for 3-5 minutes. After washing with phosphate-buffered saline (PBS) 2-3 times, worms were imaged under a fluorescence microscope.

### RNA staining

RNA was stained with SYTO™ RNASelect™ Green Fluorescent Cell Stain (SYTO™ RNASelect™ Green Fluorescent Cell Stain-5 mM Solution in DMSO, S32703, Thermo). Worms were fixed in prechilled methanol at –20°C for 10 minutes before staining and then washed twice for 5 minutes each in PBS. The labeling solution consisted of 500 nM RNA Select Green fluorescent cell stain in phosphate-buffered saline (PBS). Worms were soaked in the 500 nM labeling solution for 20 min at room temperature, washed twice in PBS for 5 min each, and then imaged.

### cRT-PCR

Total RNA was isolated from L3 stage worms using a Dounce homogenizer (pestle B) in TRIzol solution (Invitrogen). Two micrograms of total RNA was circularized by a T4 RNA Ligase 1 Kit (M0204. NEB) and then purified by TRIzol reagent followed by isopropanol precipitation. The circularized RNA was reverse transcribed via the GoScript™ Reverse Transcription System (Promega #A5001). PCR was performed using 2 × Rapid Taq Master Mix (Vazyme, P222-01) for 25 cycles. The primer sets used in this work are listed in Supplementary Table S8.

### Quantitative real-time PCR

All quantitative real-time PCR (qPCR) experiments were performed using a MyIQ2 machine (Bio-Rad). DNA or cDNA was quantified with SYBR Green Master Mix (Vazyme, Q111-02), and qPCR was performed according to the vendor’s instructions. RNA was first circularized by a T4 RNA Ligase 1 Kit (M0204. NEB) and then purified by TRIzol reagent followed by isopropanol precipitation, and then reverse transcribed via GoScript™ Reverse Transcription System (Promega #A5001) with indicated primers. The primer sets used in this work are listed in Supplementary Table S8.

### Statistics

Boxplots are presented with median and minimum and maximum. All of the experiments were conducted with independent *C. elegans* animals for the indicated N times. Statistical analysis was performed with the two-tailed Student’s t test.

## ACKNOWLEDGMENTS.

We are grateful to the members of the Guang laboratory for their comments and to Ruining Cai from the Institute of Oceanology, Chinese Academy of Sciences, Qingdao, China. We are grateful to the International C. elegans Gene Knockout Consortium and the National Bioresource Project for providing the strains. Some strains were provided by the Caenorhabditis Genetics Center (CGC), which is funded by the NIH Office of Research Infrastructure Programs (Grant P40 OD010440). This work was supported by grants from the National Key R&D Program of China (2022YFA1302700, 2019YFA0802600), the National Natural Science Foundation of China (32230016, 91940303, 32270583, 32070619, 31870812, 31871300 and 31900434), and the Strategic Priority Research Program of the Chinese Academy of Sciences (XDB39010600). This study was supported in part by the Fundamental Research Funds for the Central Universities.

## Supplemental figure legends

**Figure S1.**
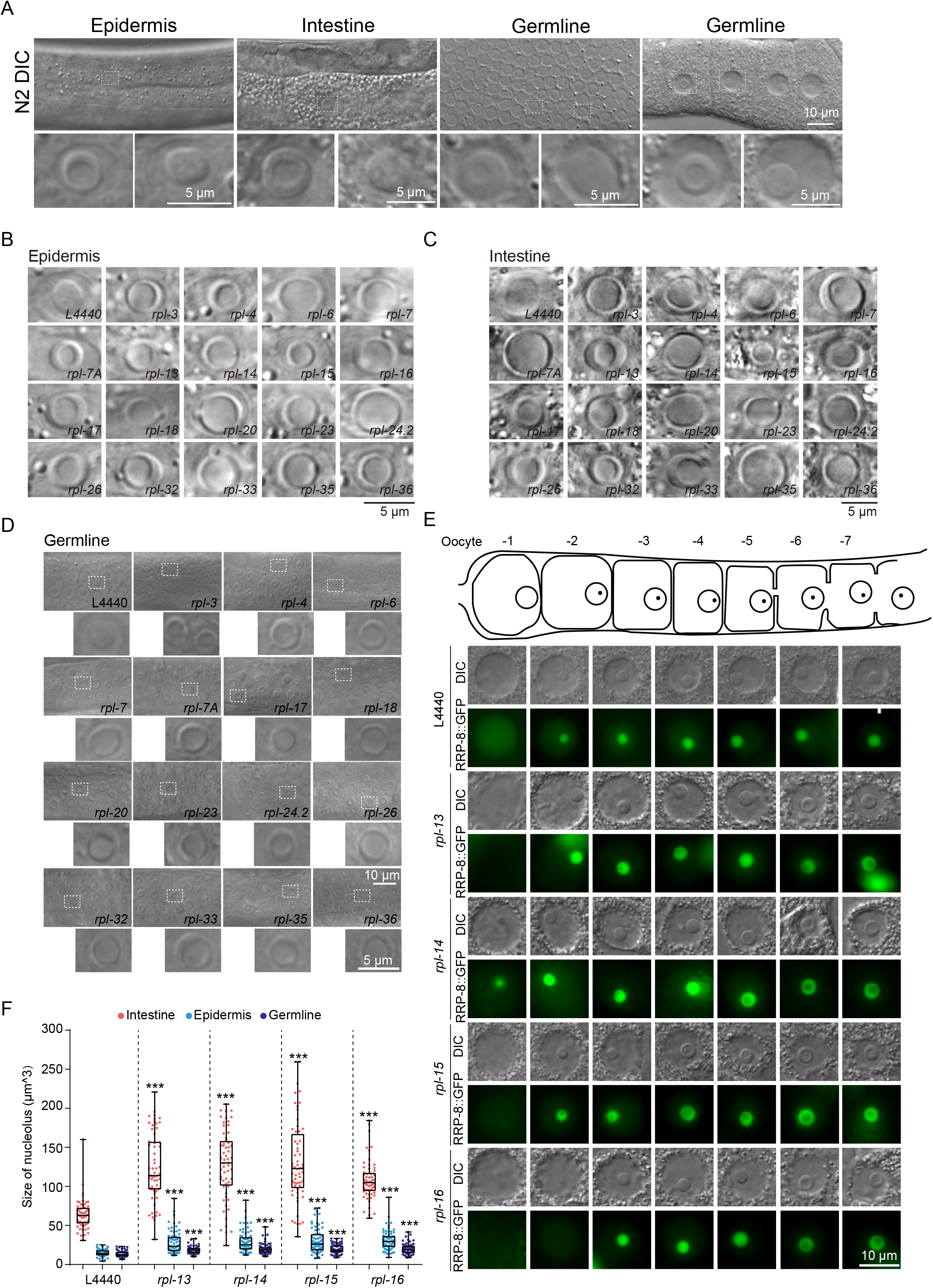
RNAi screening identified class I RPLs inhibiting the formation of ring-shaped nucleoli. (A) Differential interference contrast (DIC) of *C. elegans* nucleoli in the indicated cells. (B-D) DIC images of *C. elegans* nucleoli after knockdown of the indicated genes by RNAi. (E) Differential interference contrast (DIC) and fluorescence microscopy images of *C. elegans* nucleoli after knockdown of the indicated genes by RNAi in oocytes. (F) The size of nucleoli after RNAi knockdown of the indicated genes. n>60 nucleoli from 10 animals.

**Figure S2.**
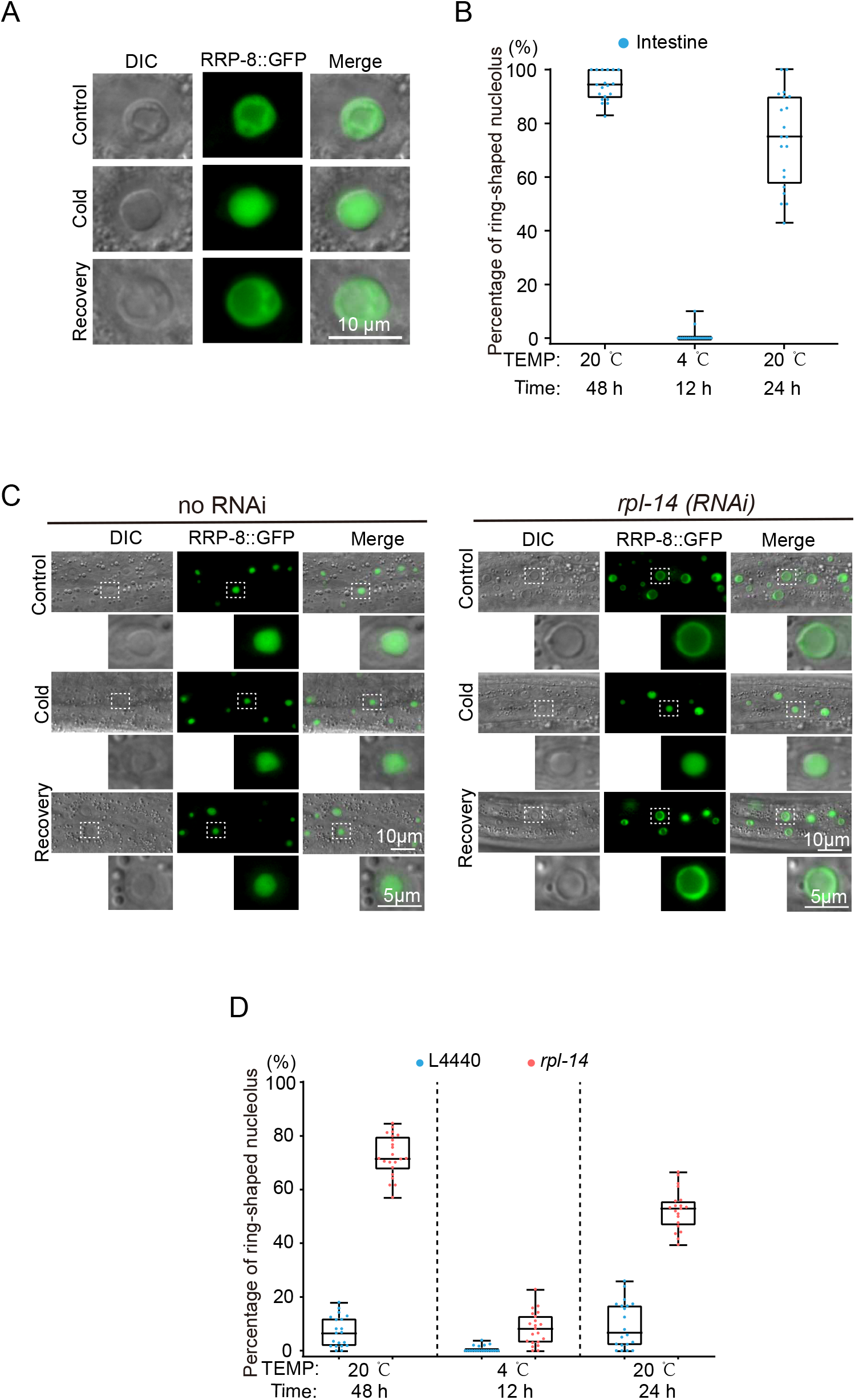
Cold stress inhibited the formation of ring-shaped nucleoli. (A-B) DIC and fluorescence microscopy images (A) and quantification (B) of ring-shaped nucleoli in intestine cells after 4°C cold stress for 12 h and recovery at 20°C for 24 h in wild-type worms. n = 20 animals. (C) DIC and fluorescence microscopy images of epidermal cells after 4°C cold stress for 12 h and recovery at 20°C for 24 h with and without *rpl-14* RNAi. (D) Quantification of ring-shaped nucleoli in epidermal cells after 4°C cold stress for 12 h and recovery at 20°C for 24 h. n = 20 animals.

**Figure S3.**
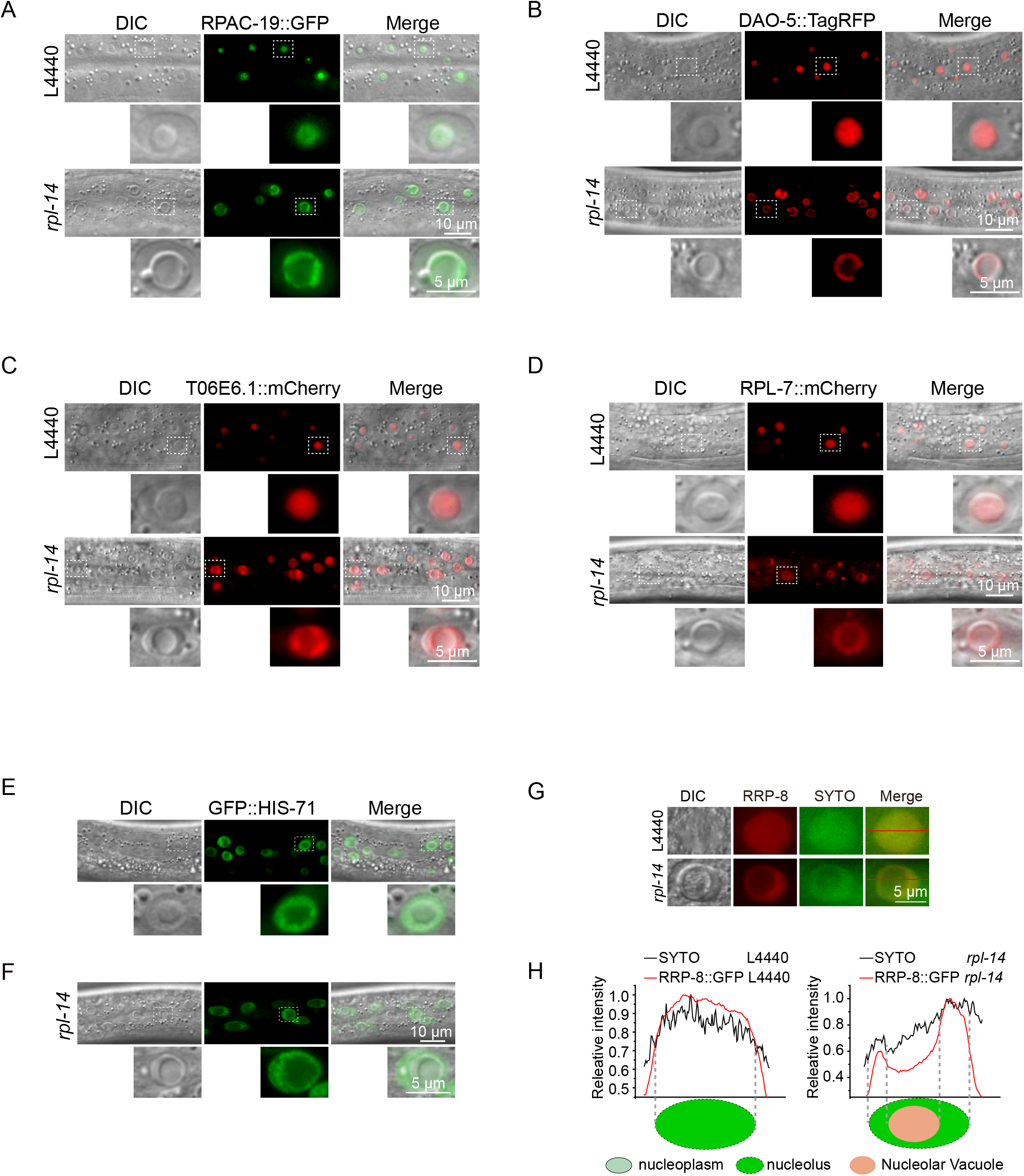
rRNA transcription and processing factors accumulate at the nucleolar ring. (A-F) DIC and fluorescence microscopy images of the indicated transgenes after knocking down *rpl-14*. (G) SYTO RNA select staining of epidermal cells after *rpl-14* RNAi. (H) Fluorescent density scan of GFP::RRP-8 and SYTO RNA select staining by ImageJ.

**Figure S4.**
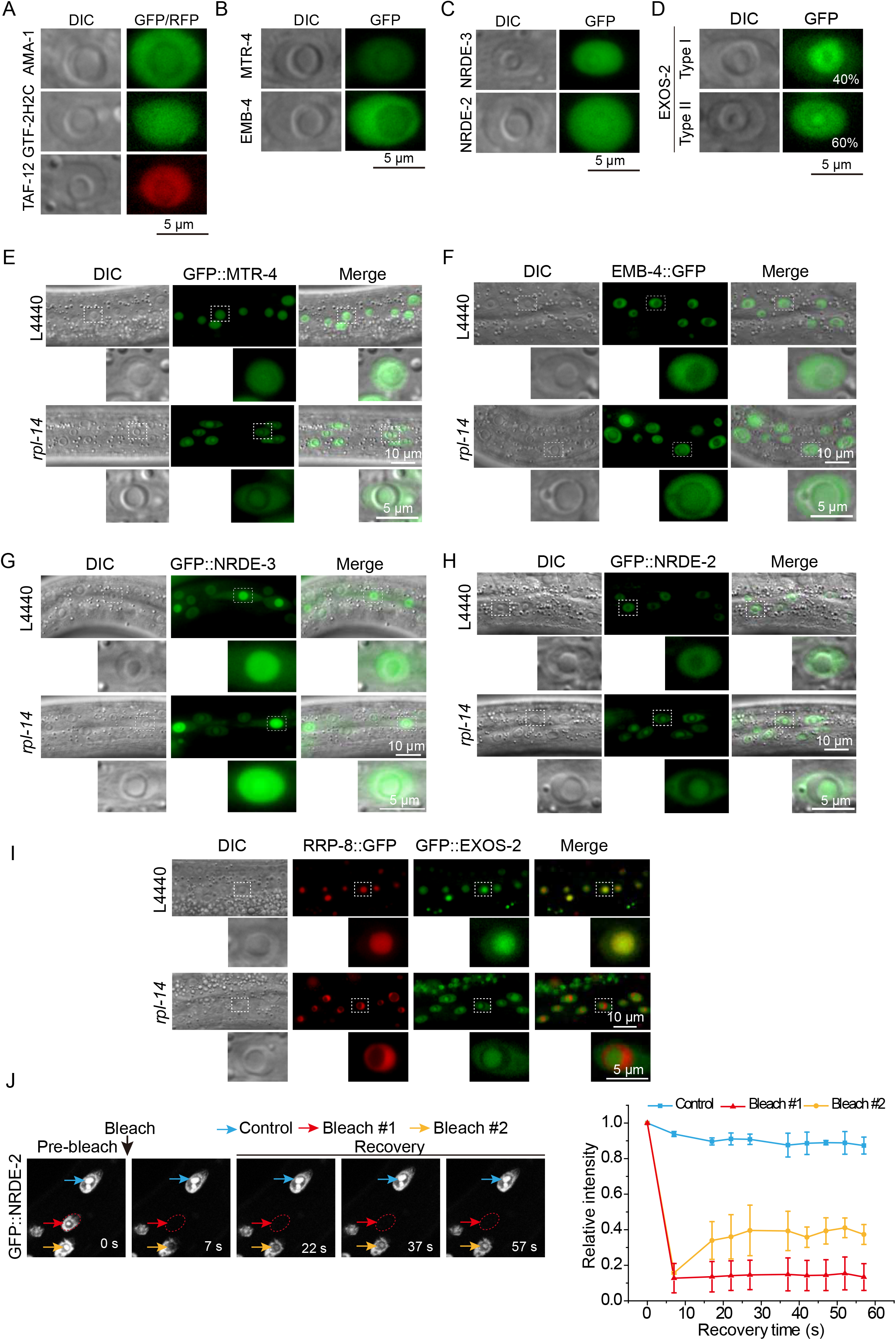
Nucleolar vacuoles contained nucleoplasmic proteins. (A-D) DIC and fluorescence microscopy images of the indicated transgenes at the ring-shaped nucleoli in wild-type animals. (E-I) DIC and fluorescence microscopy images of the indicated transgenes after knocking down *rpl-14*. (J) (left) FRAP assay of GFP::NRDE-2. (right) Quantification of FRAP data. mean ± SD, n = 3.

**Figure S5.**
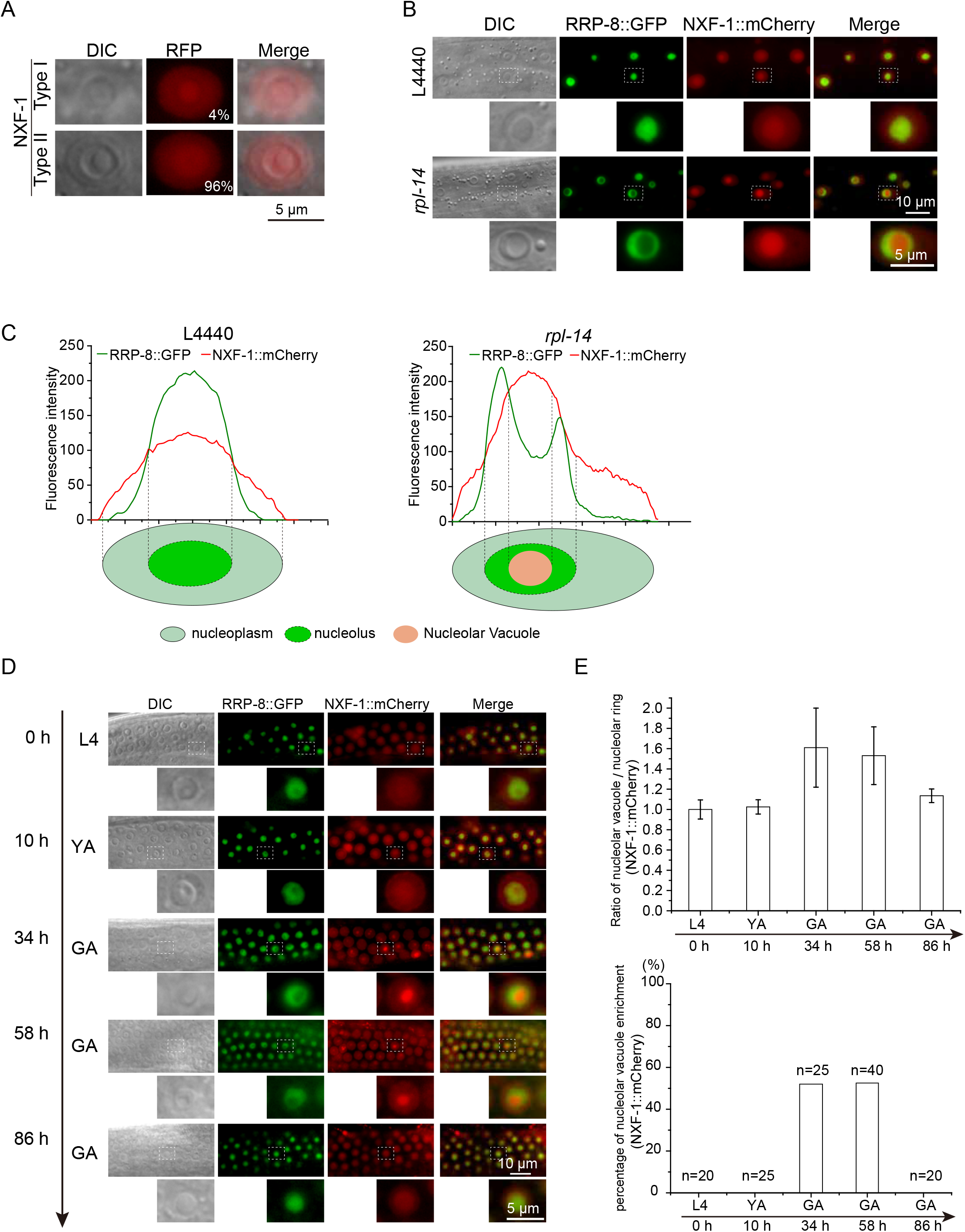
The mRNA export factor NXF-1 is enriched in the nucleolar vacuole. (A) DIC and fluorescence microscopy images of NXF-1 in ring-shaped nucleoli in wild-type animals. (B) DIC and fluorescence microscopy images of the indicated transgenes after knocking down *rpl-14*. (C) Fluorescent density scan of RRP-8::GFP and NXF-1::mCherry by ImageJ. (D) DIC and fluorescence microscopy images of the pachytene cell nucleoli in the germline. (E) (top) Relative intensity of NXF-1::mCherry in the nucleolar vacuole vs at the nucleolar ring. mean ± SD, n = 3. (bottom) Percentage of ring-shaped nucleoli at the pachytene stage. n>20 animals.

**Figure S6.**
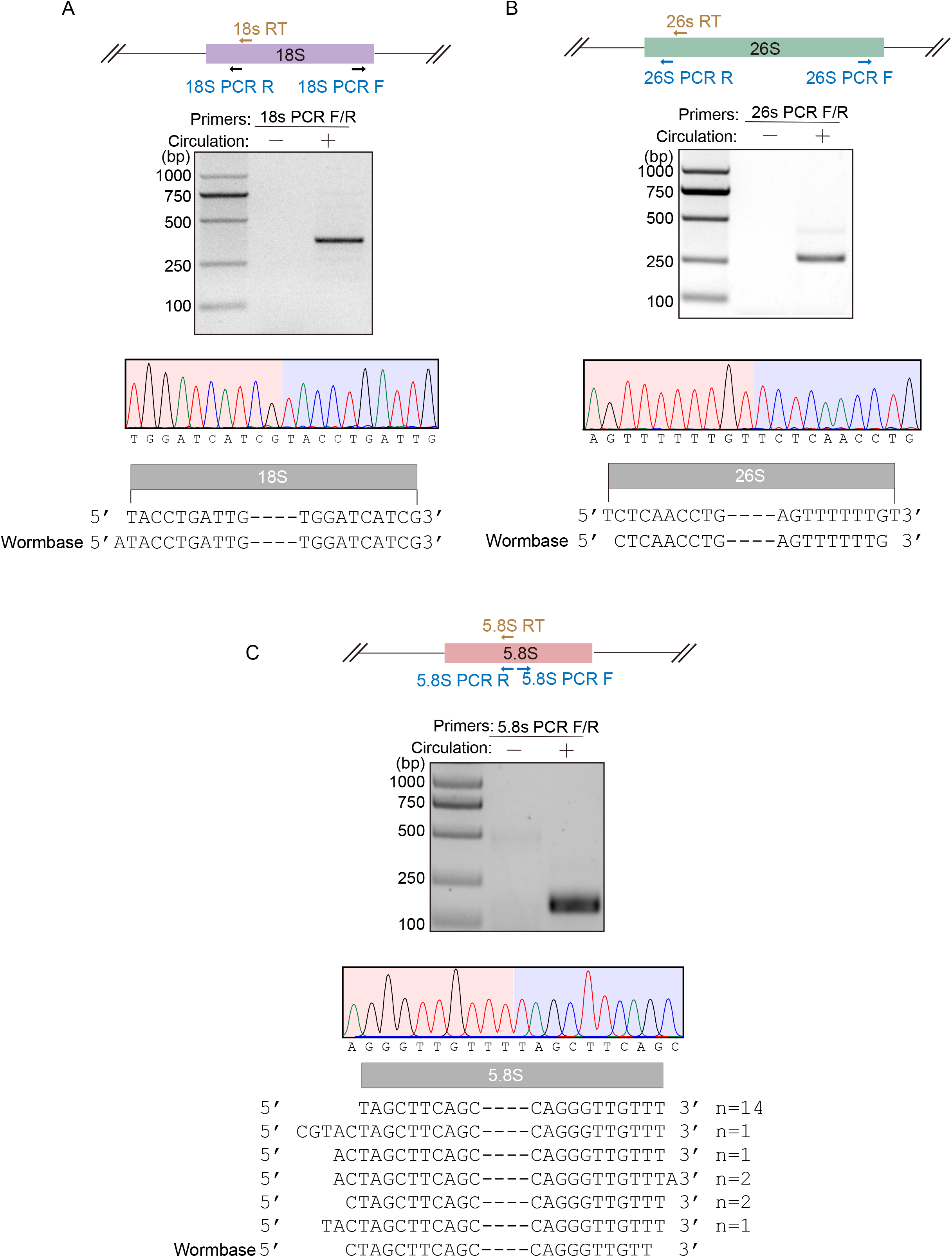
cRT-PCR assay of rRNAs. (A-C) cRT-PCR assay of mature rRNAs. Ends of the mature 18S (A), 26S (B) and 5.8S (C) rRNA were PCR amplified, followed by Sanger sequencing. Ends of mature 18S, 26S and 5.8S rRNA annotated by Wormbase are shown. N represents the number of clones sequenced.

**Figure S7.**
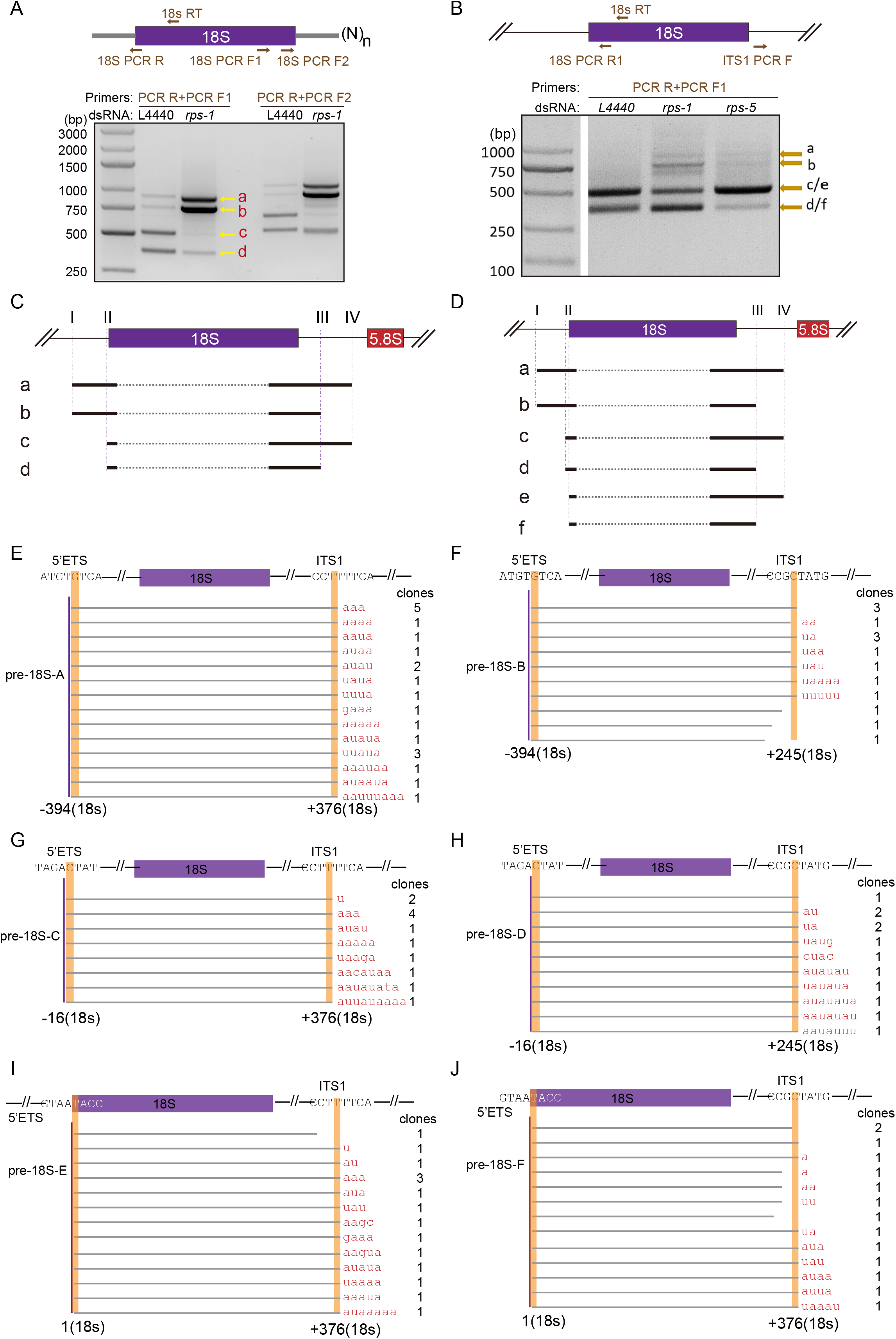
Identification of 18S rRNA intermediates by cRT-PCR. (A-B) cRT-PCR assay of 18S rRNA rRNA intermediates upon knocking down *rps-1* and *rps-5* by RNAi. (C-D) The relative positions of 18S rRNA intermediates are shown. (E-J) Sanger sequencing of the ends of six 18S rRNA intermediates. The number of clones sequenced is shown at the left of each panel.

**Figure S8.**
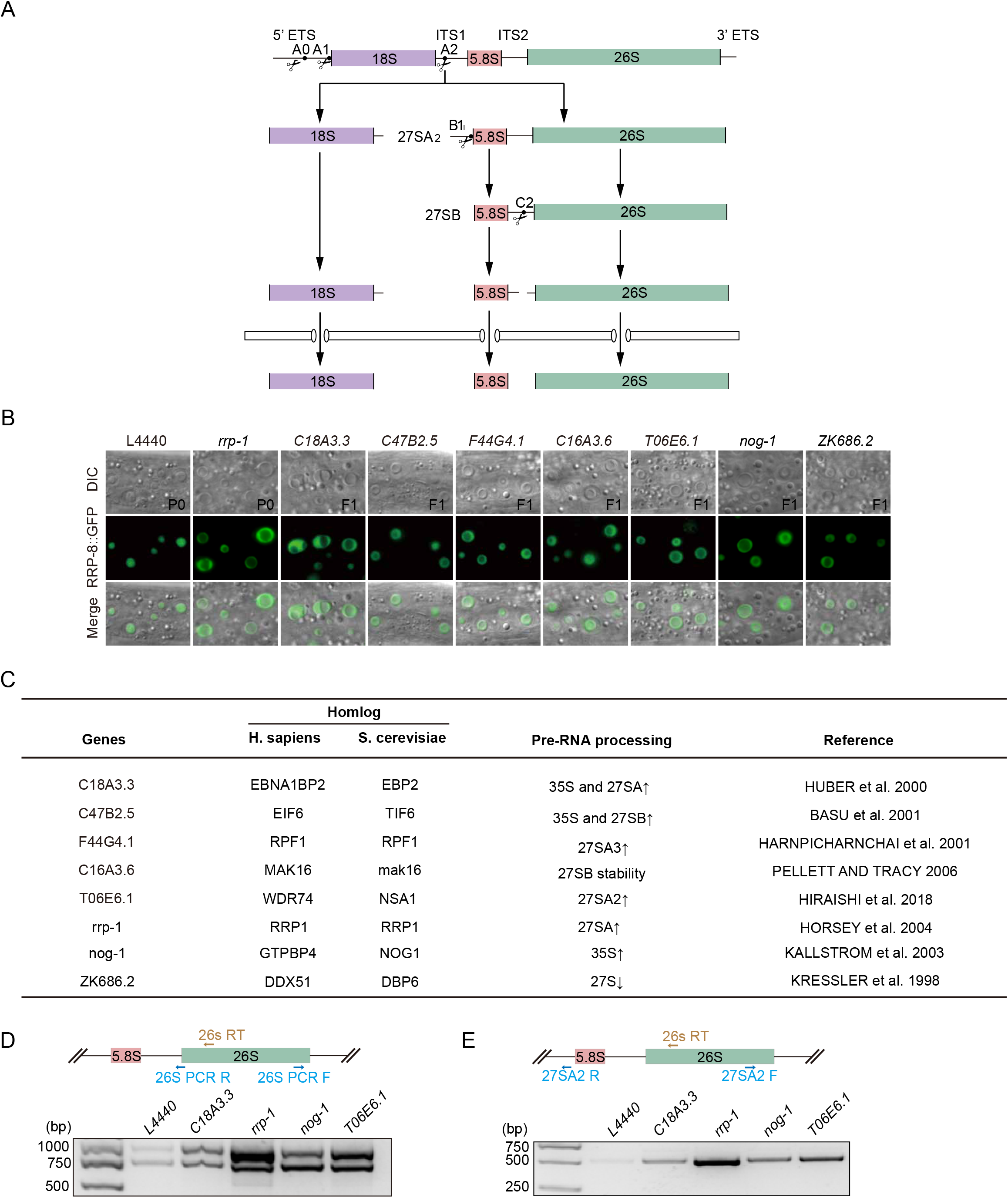
The accumulation of 27SA_2_ rRNA may disrupt nucleolar integrity. (A) Schematic diagram of rRNA processing in *C. elegans*. (B) Images of *C. elegans* nucleoli after knockdown of the indicated genes by RNAi in the epidermis. (C) Summary of the functions of the indicated genes in pre-rRNA processing. (D-E) cRT-PCR assay of 26S rRNA intermediates.

**Figure S9.**
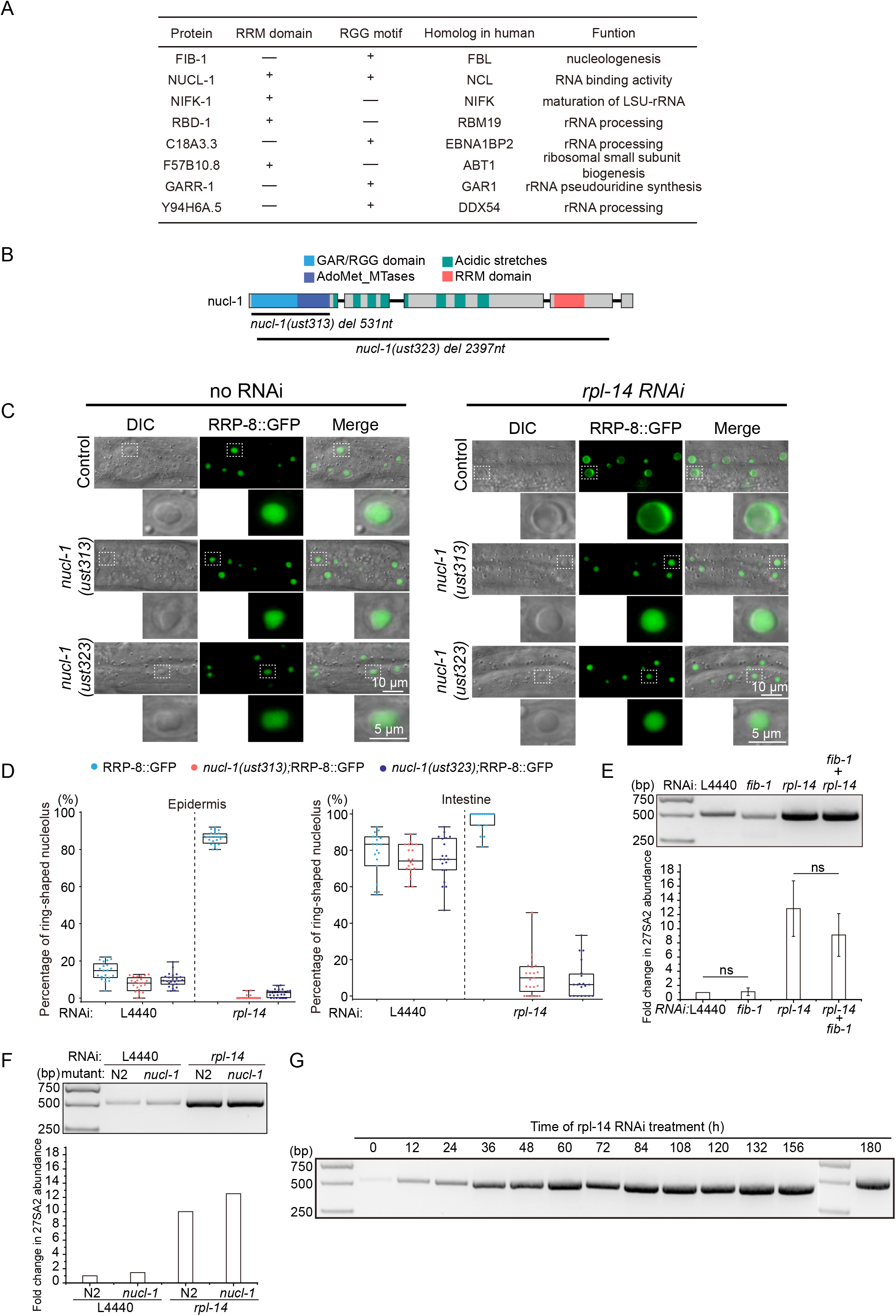
*nucl-1* and *fib-1* are required for the formation of ring-shaped nucleoli. (A) Predicated nucleolar proteins containing IDR and RNA binding motifs. (B) Schematic diagram of NUCL-1. The deletion alleles *ust313* and *ust323* were constructed by CRISPR/Cas9 technology. *ust313* was an in-frame deletion. *ust323* was likely a null allele. (C) Images of *C. elegans* nucleoli after knockdown of the indicated genes by RNAi in *nucl-1* mutants. (D) Quantification of the ring-shaped nucleoli after RNAi knockdown of the indicated genes. n>20 animals. (E) (top) cRT-PCR assay of 27SA_2_ rRNA intermediates upon knockdown of the indicated genes by RNAi. (bottom) Levels of 27SA_2_ pre-rRNAs quantified by real-time PCR. mean ± SD, n = 3. (F) (top) cRT-PCR assay of 27SA_2_ rRNA intermediates of the nucl-1 mutant upon knockdown of the indicated genes by RNAi. (bottom) Levels of 27SA_2_ pre-rRNAs quantified by real-time PCR. (G) cRT-PCR assay of 27SA_2_ rRNA intermediates by knocking down *rpl-14*.

**Table S1.** List of genes used in the candidate-based RNAi screening.

| Gene ID | Gene ID | Gene ID | Gene ID | Gene ID |
| --- | --- | --- | --- | --- |
| Y39A1A.14 | F55F8.5 | rpl-13 | rpl-33 | rps-9 |
| F18C5.3 | C15H11.9 | rpl-14 | rpl-34 | rps-10 |
| ZK1127.5 | W09C5.1 | rpl-15 | rpl-35 | rps-11 |
| T22H9.1 | T19A6.2 | rpl-16 | rpl-36 | rps-12 |
| ZC434.4 | K01C8.9 | rpl-17 | rpl-36A | rps-13 |
| T22H6.1 | W07E6.2 | rpl-18 | rpl-37 | rps-14 |
| Y54E5B.2 | T25G3.3 | rpl-19 | rpl-38 | rps-15 |
| Y75B8A.7 | Y56A3A.18 | rpl-20 | rpl-39 | rps-16 |
| C48B6.2 | crn-3 | rpl-21 | rpl-40 | rps-17 |
| F58B3.4 | rpl-1 | rpl-22 | rpl-41.1 | rps-18 |
| F57B10.8 | rpl-2 | rpl-23 | rpl-41.2 | rps-19 |
| ZK430.7 | rpl-3 | rpl-24.1 | rpl-42 | rps-20 |
| T20B12.3 | rpl-4 | rpl-24.2 | rpl-43 | rps-21 |
| F57B9.5 | rpl-5 | rpl-25.1 | rps-0 | rps-22 |
| Y48B6A.3 | rpl-6 | rpl-25.2 | rps-1 | rps-23 |
| T01C3.7 | rpl-7 | rpl-26 | rps-2 | rps-24 |
| K07C5.4 | rpl-7A | rpl-27 | rps-3 | rps-25 |
| Y6B3B.9 | rpl-9 | rpl-28 | rps-4 | rps-26 |
| F54C1.2 | rpl-10 | rpl-29 | rps-5 | rps-27 |
| rrp-8 | rpl-11.1 | rpl-30 | rps-6 | rps-28 |
| R13A5.12 | rpl-11.2 | rpl-31 | rps-7 | rps-29 |
| F23B12.7 | rpl-12 | rpl-32 | rps-8 | rps-30 |

**Table S2.**
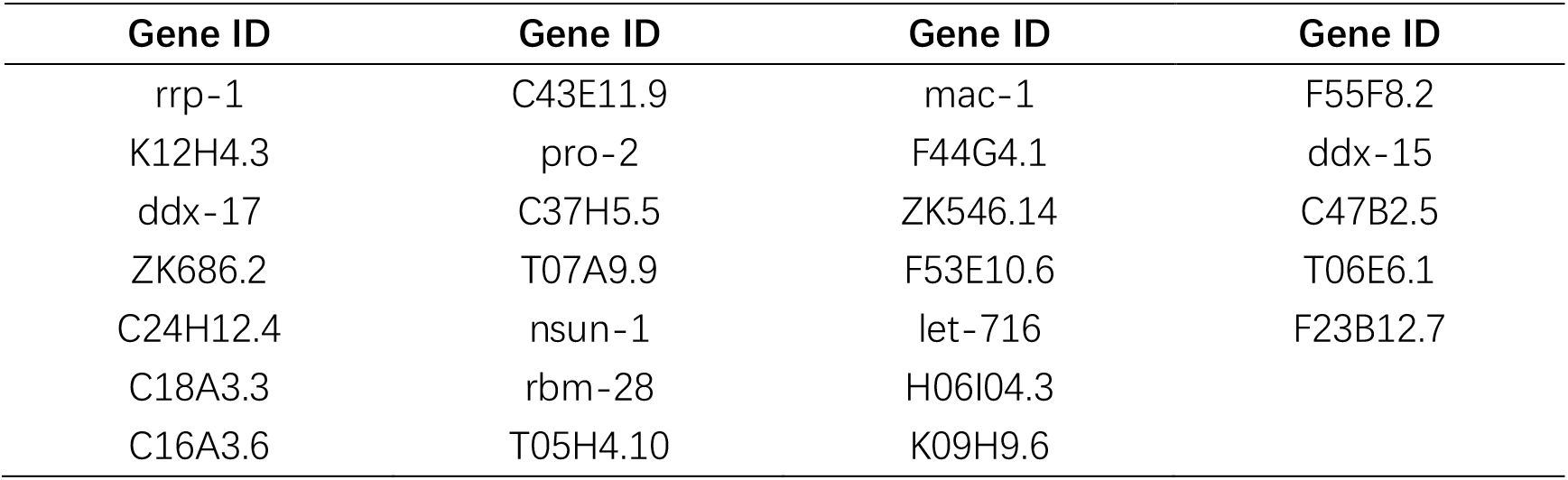
List of genes involved in 26S rRNA processing.

| **Gene ID** | **Gene ID** | **Gene ID** | **Gene ID** |
| --- | --- | --- | --- |
| rrp-1 | C43E11.9 | mac-1 | F55F8.2 |
| K12H4.3 | pro-2 | F44G4.1 | ddx-15 |
| ddx-17 | C37H5.5 | ZK546.14 | C47B2.5 |
| ZK686.2 | T07A9.9 | F53E10.6 | T06E6.1 |
| C24H12.4 | nsun-1 | let-716 | F23B12.7 |
| C18A3.3 | rbm-28 | H06I04.3 |  |
| C16A3.6 | T05H4.10 | K09H9.6 |  |

**Table S3:** List of strains used in this study.

| Strains | Genotype |
| --- | --- |
| N2 |  |
| shg2123 | rrp-8::gfp::3xflag (ustIS76) |
| shg2287 | fib-1P::fib-1::mCherry::rps-1 3' UTR (ustIS293) |
| shg1248 | 3xflag::gfp::rpoa-2 (ustIS116) |
| shg2285 | rpl-7::mCherry (ustIS291) |
| shg1092 | rrp-8::mCherry (ustIS128) |
| shg1268 | 3xflag::gfp::rpoa-1 (ustIS137) |
| shg1229 | 3xflag::gfp::rpac-19 |
| shg1889 | tagRFP::dao-5 (ustIS242) |
| shg1660 | rbd-1::mCherry (ustIS207) |
| shg2286 | T06E6.1::mCherry (ustIS292) |
| shg852 | 3xflag::gfp::nrde-2 (ustIS117) |
| shg664 | mtr-4p::gfp::mtr-4::mtr-4 UTR |
| shg2288 | ama-1::gfp::3xflag (ust423) |
| YY174 | 3xflag::gfp::nrde-3 (ggIS1) |
| shg011 | *eri-1(mg366)*; ggIS1;susi-1(ust001) |
| shg1890 | 3xflag::gfp::his-71 (ustIS254) |
| shg2160 | 3xflag::gfp::gtf-2H2C (ust384) |
| shg1696 | taf-12::tagRFP (ustIS227) |
| shg2289 | emb-4::gfp::3xflag (ust424) |
| shg2141 | nxf-1::mCherry (ustIS281) |
| shg2142 | nxf-1::mCherry(ustIS281);rrp-8::gfp(ustIS76) |
| shg2104 | 3xflag::gfp::exos-2(ustIS113);rrp-8::mCherry (ustIS128) |
| shg1914 | *nucl-1 (ust313)* |
| shg2090 | *nucl-1 (ust323)* |
| shg2098 | *nucl-1 (ust313)*; susi-2::gfp(ustIS76) |
| shg2143 | *nucl-1 (ust323)*; susi-2::gfp(ustIS76) |

**Table S4.**
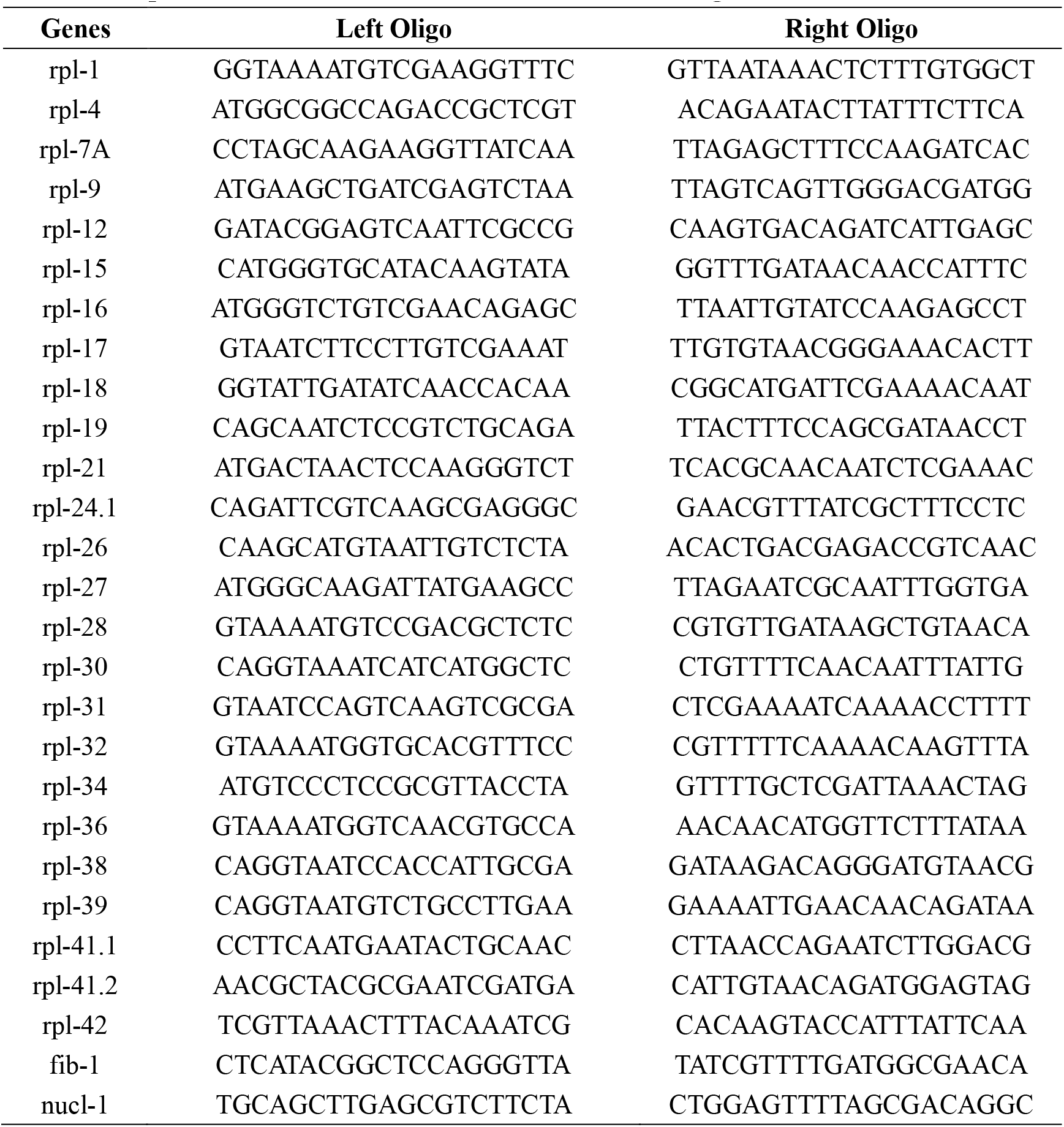
Sequences of double-stranded RNA for RNAi screening.

**Table S5.**
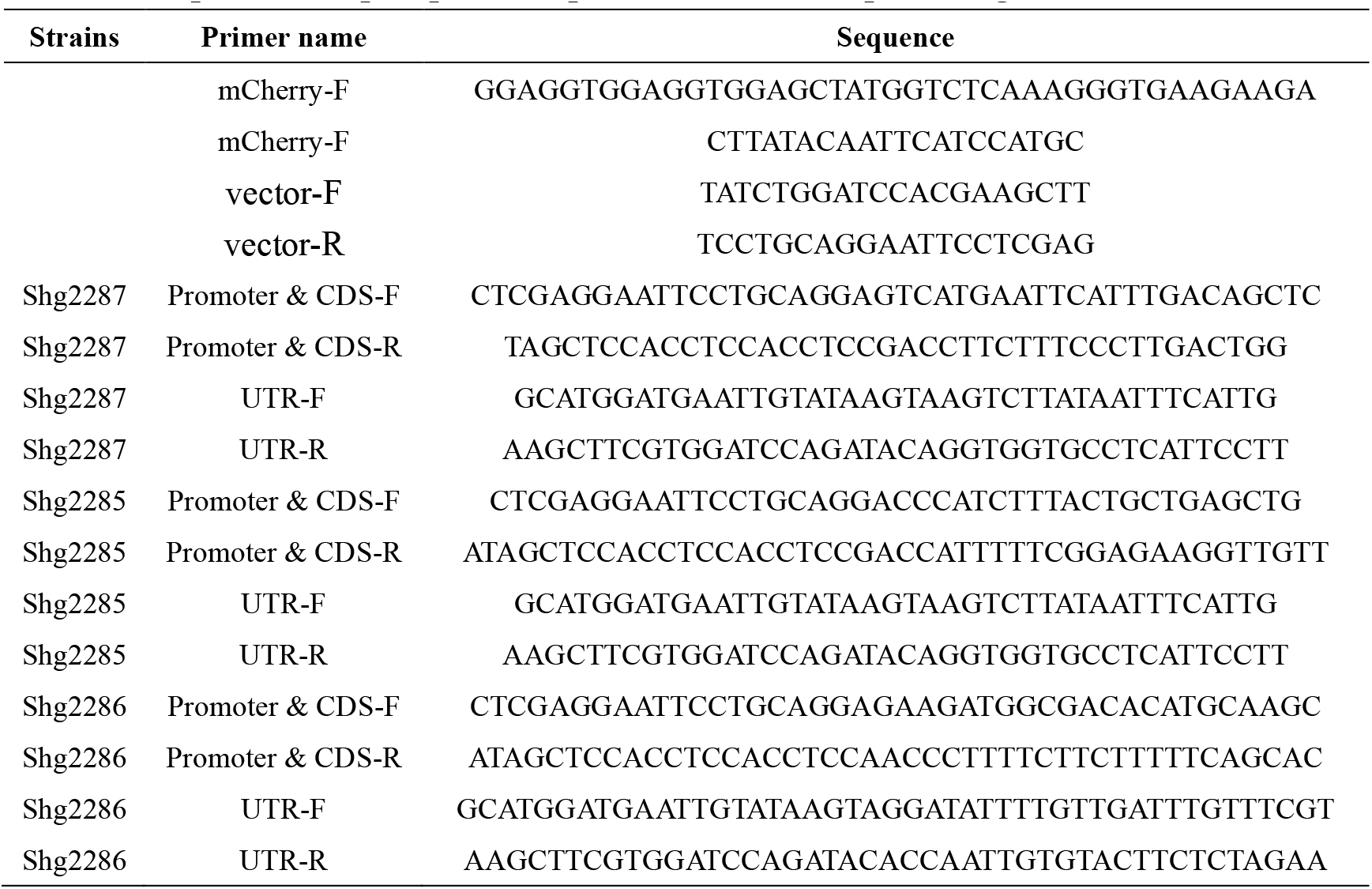
Sequence of repair plasmid primers used in ectopic transgenic strain construction.

**Table S6.**
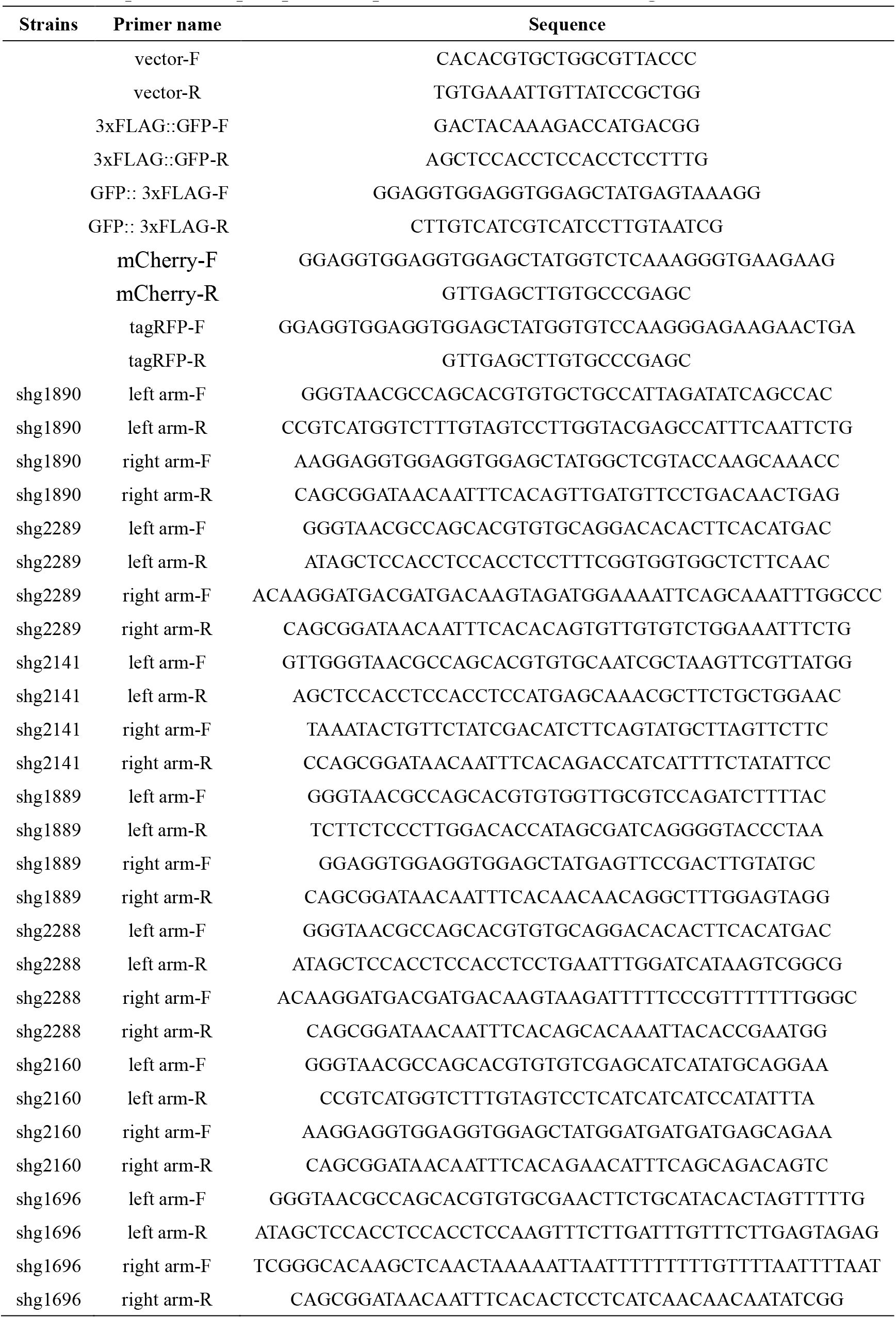
Sequence of repair plasmid primers used in in situ transgenic strain construction.

**Table S7.**
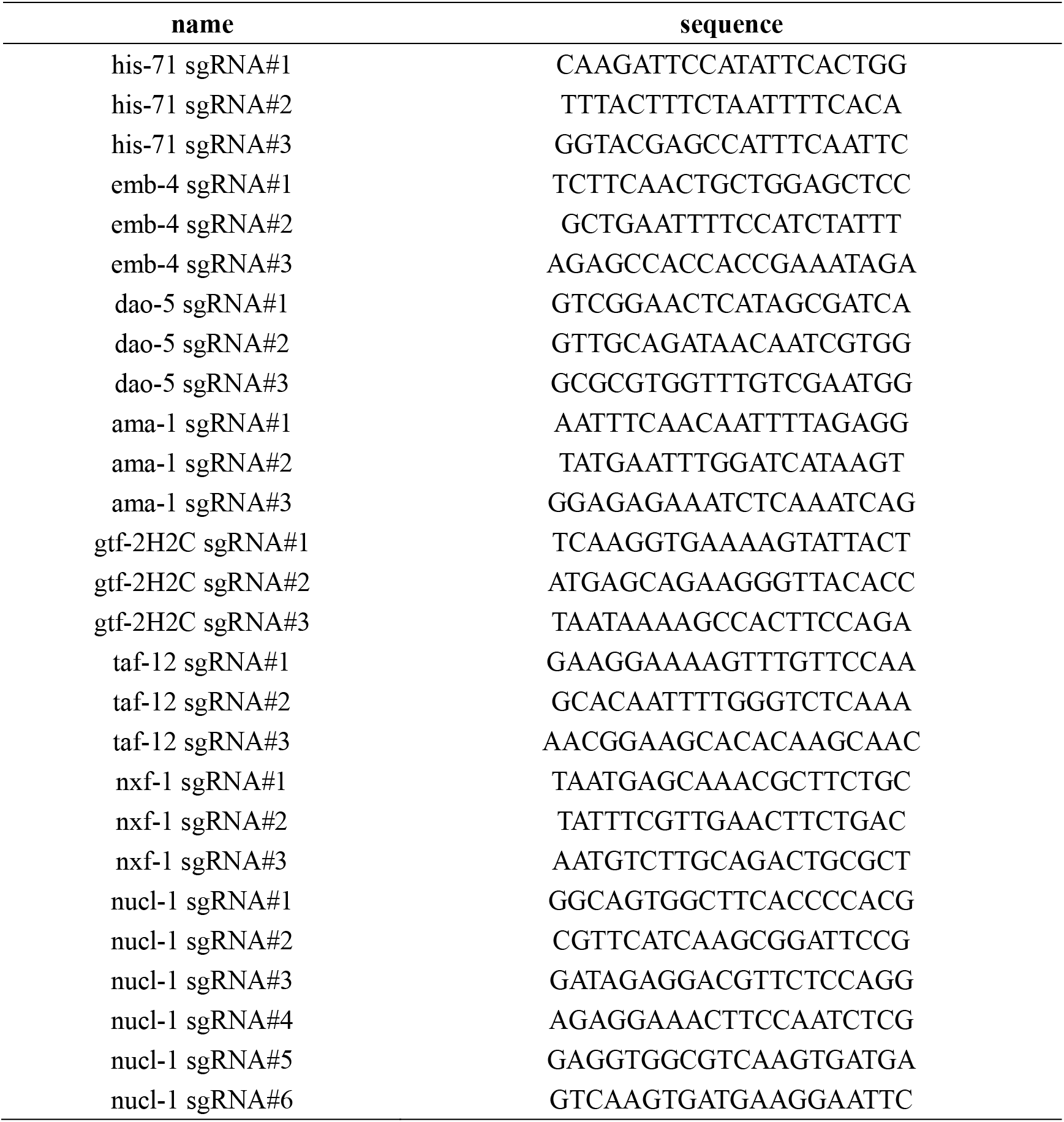
Sequences of sgRNAs for CRISPR/Cas9-mediated gene editing.

**Table S8.**
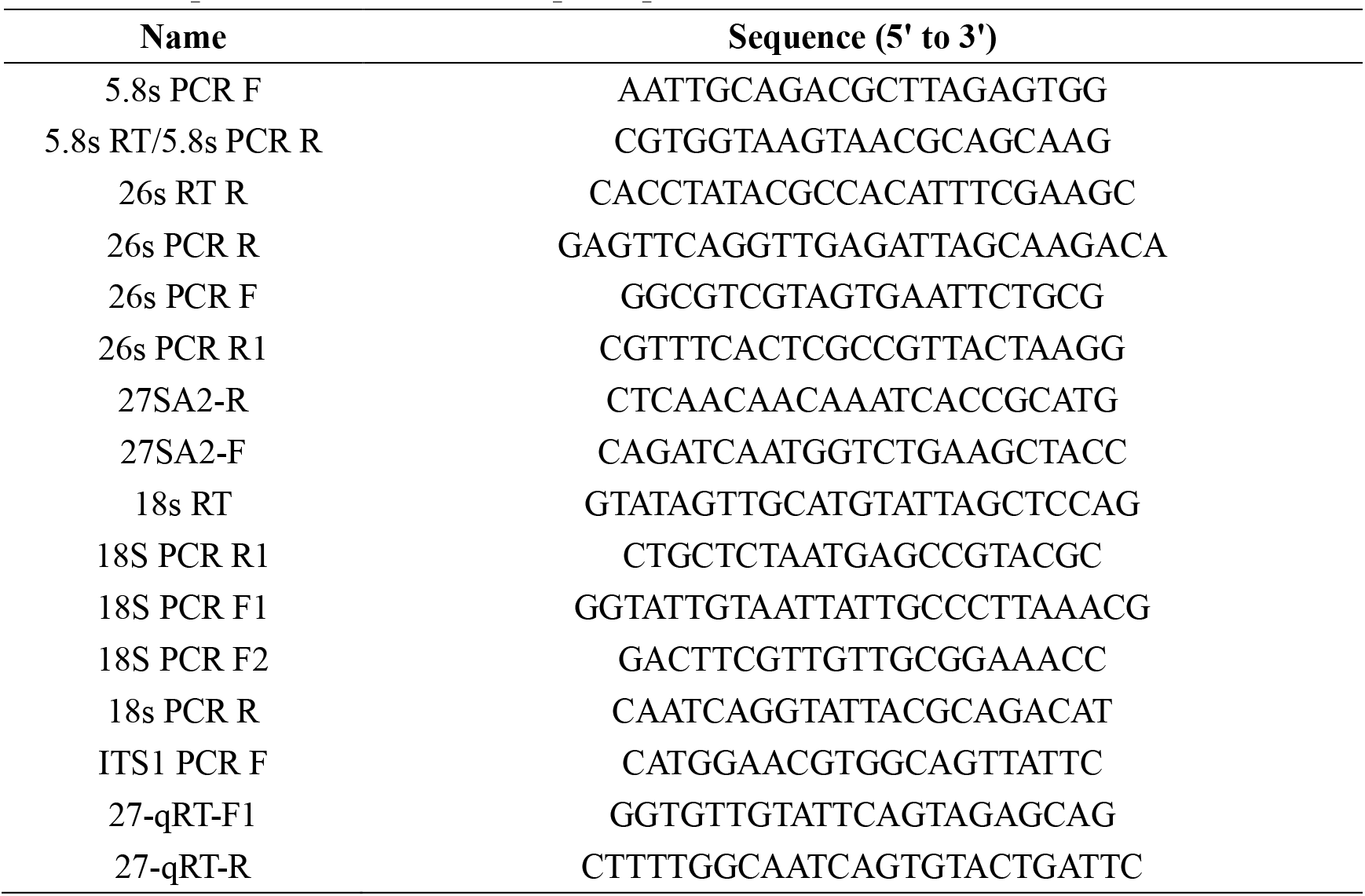
Sequence of cRT-PCR and qPCR primers.

## Notes

### Competing Interest Statement

The authors have declared no competing interest.

